# Transgenic expression of the dicotyledonous pattern recognition receptor EFR in rice leads to ligand-dependent activation of defense responses

**DOI:** 10.1101/006155

**Authors:** Benjamin Schwessinger, Ofir Bahar, Nicolas Thomas, Nicolas Holton, Vladimir Nekrasov, Deling Ruan, Patrick E. Canlas, Arsalan Daudi, Christopher J. Petzold, Vasanth R Singan, Rita Kuo, Mansi Chovatia, Christopher Daum, Joshua L. Heazlewood, Cyril Zipfel, Pamela C. Ronald

## Abstract

Plant plasma membrane localized pattern recognition receptors (PRRs) detect extracellular pathogen-associated molecules. PRRs such as Arabidopsis EFR and rice XA21 are taxonomically restricted and are absent from most plant genomes. Here we show that rice plants expressing EFR or the chimeric receptor EFR::XA21, containing the EFR ectodomain and the XA21 intracellular domain, sense both *Escherichia coli*- and *Xanthomonas oryzae* pv. *oryzae* (*Xoo*)-derived elf18 peptides at sub-nanomolar concentrations. Treatment of EFR and EFR::XA21 rice leaf tissue with elf18 leads to MAP kinase activation, reactive oxygen production and defense gene expression. Although expression of EFR does not lead to robust enhanced resistance to fully virulent Xoo isolates, it does lead to quantitatively enhanced resistance to weakly virulent Xoo isolates. EFR interacts with OsSERK2 and the XA21 binding protein 24 (XB24), two key components of the rice XA21-mediated immune response. Rice-EFR plants silenced for *OsSERK2*, or overexpressing rice *XB24* are compromised in elf18-induced reactive oxygen production and defense gene expression indicating that these proteins are also important for EFR-mediated signaling in transgenic rice. Taken together, our results demonstrate the potential feasibility of enhancing disease resistance in rice and possibly other monocotyledonous crop species by expression of dicotyledonous PRRs. Our results also suggest that Arabidopsis EFR utilizes at least a subset of the known endogenous rice XA21 signaling components.

**Author Summary:** Plants possess multi-layered immune recognition systems. Early in the infection process, plants use receptor proteins to recognize pathogen molecules. Some of these receptors are present in only in a subset of plant species. Transfer of these taxonomically restricted immune receptors between plant species by genetic engineering is a promising approach for boosting the plant immune system. Here we show the successful transfer of an immune receptor from a species in the mustard family, called EFR, to rice. Rice plants expressing EFR are able to sense the bacterial ligand of EFR and elicit an immune response. We show that the EFR receptor is able to use components of the rice immune signaling pathway for its function. Under laboratory conditions, this leads to an enhanced resistance response to two weakly virulent isolates of an economically important bacterial disease of rice.

## Introduction

Plants possess multi-layered immune systems enabling them to fend off most pathogens. Plasma membrane localized pattern recognition receptors (PRRs) sense danger-associated molecules, including pathogen/microbe-associated molecular patterns (P/MAMPs) and endogenous elicitors released during the infections process. The activation of PRRs triggers a rapid intracellular signaling cascade [1–3]. In most cases, PRR-triggered immunity (PTI) is sufficient to halt microbial replication and disease development 4,5]. Successful pathogens are able to suppress PTI by employing effector molecules in the apoplast or inside the plant cell and thereby enable plant colonization. Plants, in turn, recognize specific effector molecules either in the apoplast via transmembrane receptors or inside the cell via intracellular immune receptors, which are part of the nucleotide binding site-leucine rich repeat (NBS-LRR) protein family. This recognition event is often referred to as effector triggered immunity (ETI) [6,7,4,8].

An important goal of plant pathology research is to generate knowledge that can be used to enhance resistance to serious diseases in crops [9]. An emerging approach that has recently been successful is the transfer of plasma membrane-localized PRRs between distinct plant species [10–14]. All well—defined plant PRRs are receptor kinases (RKs) or receptor-like proteins (RLPs) [1,2,15,16]. This class of immune receptors senses conserved microbial molecules such as bacterial flagellin, bacterial elongation factor Tu (EF-Tu) and fungal chitin [1,2,15]. Flagellin and its corresponding receptor FLAGELLIN SENSING 2 (FLS2; At5G6330) are the best-studied ligand-plant PRR pair [2]. In many plant species, FLS2 recognizes a conserved 22-amino-acid-long internal epitope originally derived from flagellin of *Pseudomonas syringae* pv. *tabaci* (flg22_*Pta*_) [17–19]. Other plant species are able to recognize additional epitopes of flagellin [20–25].

In contrast to the wide conservation of the flagellin-FLS2 perception system, other PRRs and their respective PAMP recognition specificity are restricted to limited plant families or species [1,3,15]. For example, EF-TU RECEPTOR (EFR; At5g20480), the receptor that recognizes the highly conserved N-terminal 18 amino acids of EF-Tu, originally isolated from *Escherichia coli* (from here on referred to as elf18_*E.coli*_), is restricted to the plant family *Brassicaceae* [26,27]. Similarly, XA21 (U37133), which recognizes the bacterial pathogen *Xanthomonas oryzae* pv. *oryzae (Xoo)*, appears to be restricted to the wild rice species *Oryza longistaminata* [28–30]. These taxonomically restricted PRRs are prime candidates for inter-species transfer, because they may be able to confer resistance to a wide range of pathogens for which there is presently no disease control measures. Indeed, it was recently shown that the inter-family transfer of Arabidopsis EFR to tomato and *Nicotiana benthamiana*, both members of the *Solanaceae* family, confers the plants with the ability to recognize elf18. Furthermore, tobacco and tomato plants expressing EFR became more resistant to a phylogenetically diverse range of bacterial pathogens including *P. syringae* pv. *tabaci* and *R. solanacearum*, respectively [10]. Also the transfer of rice XA21 into citrus *(Citrus sinensis)*, tomato (*Lycopersicon esculentum*) and banana (*Musa sp*.) confers moderate resistance to *X. axonopodis* pv. *citri* and strong resistance to *Ralstonia solanacearum* and X. *campestris* pv. *malvacearum*, respectively [11–13]. Similarly, the inter-family transfer of the tomato RLP Ve1 (NM_001247545.1), which recognizes Ave1 from *Verticillium dahlia* race1 [31], to Arabidopsis confers race-specific resistance to *V. dahlia* [14]. These transfers of taxonomically restricted PRRs between different dicot species or from a monocot to a dicot species demonstrate that this strategy is a viable approach to improve plant immunity at least under controlled conditions. No field tests have been performed with crops transgenically expressing taxonomically restricted PRRs, which are necessary for full evaluation of the effectiveness of this engineered resistance strategy.

It is not yet known if the transfer of a dicotyledonous PRR, such as EFR, into an important monocotyledonous staple crops species, such as rice, provides resistance to bacterial infection. Assessment of the functionality of this directional inter-class transfer of PRRs is of broad interest because monocotyledonous cereals such as rice, wheat, and corn generate 80% of the calories consumed by humans according to the Food and Agriculture Organization of the United Nations. In addition, most of the molecular knowledge of plant immune receptors, including PRRs, has been gained from studies of dicotyledonous model systems [1,2,15].

Many of the genetic and biochemical requirements for downstream signaling of EFR in Arabidopsis have been characterized. Before EFR reaches the plasma membrane, it undergoes substantial folding and post-translation glycosylation in the endoplasmatic reticulum (ER) and Golgi apparatus. These modifications are important for ligand binding and proper function [32–36]. At the plasma membrane, EFR forms heteromeric complexes with at least four coreceptor-like kinases belonging to the SOMATIC EMBRYOGENESIS RECEPTOR KINASE (SERK) family within seconds to minutes of ligand binding [37–42]. Ligand perception induces rapid phosphorylation of EFR including tyrosine phosphorylation, which is important for full downstream signal activation [38,43]. The interaction between EFR and SERK proteins leads to the activation and release of BOTRYTIS INDUCED KINASE 1 (BIK1; At2g39660) and additional members of the cytoplasmic receptor-like kinase subfamily VII from the complex [44,45]. Further EFR downstream signaling events involves a Ca^2+^ influx, reactive oxygen species (ROS) production via NADPH oxidases at the plasma membrane and apoplastic peroxidases, Ca^2+^-dependent kinases and mitogen-activated protein kinase (MAPK) cascades [2,46–49]. These partially independent signaling cascades cumulate in significant transcriptional reprogramming involving several WRKY transcription factors [26,50].

Similarly to EFR, XA21 biogenesis occurs in the ER [51,52]. After processing and transit to the plasma membrane, XA21 binds to XB24 (XA21 Binding Protein 24, Os01g56470) [53]. XB24 is a rice specific ATPase that binds to the XA21 juxtamembrane domain and uses ATP to promote phosphorylation of certain Ser/Thr sites on XA21, keeping the XA21 protein in an inactive state. Upon recognition of *Xoo*, the XA21 kinase disassociates from XB24 and is activated, triggering downstream defense responses [53]. XA21 interacts constitutively with OsSERK2 (Os04g38480) and requires OsSERK2 for full downstream signaling initiation [54]. Key components of the downstream response include MAPKs [55], a RING finger ubiquitin ligase (XB3, AF272860) [56], the plant-specific ankyrin-repeat (PANK) containing protein XB25 (Os09g33810) [57], and WRKY transcription factors OsWRKY62 and 76 (NP_001063185 and DAA05141) [58]. XA21 activity is down-regulated by dephosphorylation post-defense activation by the protein phosphatase 2C XB15 (Os03g60650) [59].

EFR and XA21 are phylogenetically closely related (Supplementary Figure 1) [60] and share some, but not all, orthologous signaling components. For example, both receptor require the ER quality machinery for proper folding and function [32–36,51,52]. Orthologous SERK family members interact with both receptors and are required for downstream signaling initiation in both Arabidopsis and rice [41,42,54]. In both cases, WRKY transcription factors are involved in transcriptional reprogramming [50,58]. However, it is not yet known if EFR activity in Arabidopsis is down regulated by dephosphorylation, if an ATPase is required for its inactivation in the absence of the ligand or if an E3 ligase is important for its stability. It is therefore very difficult to predict if EFR would be functional when transgenically expressed in rice and if it would employ the same signaling components as the endogenous rice PRR XA21.

Here we report that the expression of EFR or the chimeric receptor EFR::XA21 makes rice receptive to elf18_*Ecoli*_ at sub-nanomolar concentrations, inducing MAP kinase activation, ROS production, and defense gene expression. We show that EF-Tu is highly conserved in over twenty different *Xoo* isolates and that elf18_*Xoo*_, which carries two amino acid substitutions at position 2 and 4 in comparison with elf18_*E.coli*_, is fully recognized by rice plants expressing EFR at similar sub-nanomolar concentrations. The recognition of elf18_*Xoo*_ in rice plants expressing EFR leads to moderate quantitatively enhanced resistance to weakly virulent isolates of Xoo. In contrast, the EFR-rice plants were only slightly more resistant to fully virulent isolates of *Xoo* in 3 out of 6 experiments. Surprisingly, rice-EFR::XA21 plants did not become fully resistant to *Xoo* instead displaying a weak resistance profile similar to rice-EFR plants. We further demonstrate that EFR directly interacts with two signaling components of XA21, OsSERK2 and XB24, and both are required for EFR signaling in rice.

## Results

### Expression of EFR or the chimeric receptor EFR::XA21 in rice confers responsiveness to elf18_*E.coli*_

We generated two constructs to test if the expression of the EFR ectodomain enables rice to sense and respond to elf18_*E.coli*_. The first construct expresses full-length *Arabidopsis* EFR with a carboxyl-terminal green fluorescent protein (GFP) fusion under the control of the maize ubiquitin promoter (*Ubi::EFR::GFP*) [26]. The second construct expresses the EFR ectodomain (EFR 1-649 aa) fused to the XA21 transmembrane, juxtamembrane and intracellular domains (XA21 651-1025 aa) with a carboxyl-terminal GFP fusion under the control of the maize ubiquitin promoter *(Ubi::EFR::XA21::GFP)* (Supplementary Figure 2) [26,28]. We reasoned that the XA21 kinase domain might be more adapted to intracellular signal initiation in rice. We obtained six independent PCR positive T_0_ rice transformants for *Ubi::EFR::GFP* and five in the case of *Ubi::EFR::XA21::GFP.* Four of the T_0_ lines for *Ubi::EFR::GFP* (−4, −6, −7 and −9) and three of the T_0_ lines for *Ubi::EFR::XA21::GFP* (−1, −3 and −4) expressed detectable full length protein in the T_1_ generation (Supplementary Figure 3). The molecular weight of both GFP fusion proteins was similar to that of XA21::GFP of ~175kDa. This is well above the predicted molecular weight of ~140kDa and suggests that the ectodomain of EFR undergoes post-translation modification when expressed in rice. This is similar to the observation made in Arabidopsis where EFR also migrates well above its predicted molecular weight due to the glycosylation of its extracellular domain. It was previously shown that proper glycosylation of EFR is essential for its function [26,32,34–36]. We next chose three PCR-positive lines for each construct (T_0_ lines *Ubi::EFR::GFP* −1, −7, −9 and *Ubi::EFR::XA21::GFP* −2, −3, −4) and performed in depth functional analyses of the T_1_ progeny. Based on the transgene segregation analysis in the T_1_, all T_0_ lines carry a single T-DNA insertion (Supplementary Table 1). First, we assessed the transcript level of *EFR::GFP* or *EFR::XA21::GFP* with primers annealing to the sequence corresponding to the EFR ectodomain. As shown in Figure 1 A, all lines except for *Ubi::EFR::GFP*-1 expressed the transgene to comparable levels. We also tested the expression of both constructs at the protein level. Consistent with the qRT-PCR results, we were not able to detect any protein in line *Ubi::EFR::GFP-*1. All other lines expressed full-length EFR::GFP or EFR::XA21::GFP (Figure 1 B, Supplementary Figure 4). To assess if the expression of EFR::GFP or EFR::XA21::GFP enables rice to sense and respond to elf18_*E.coli*_, we measured the expression of two well-established rice defense marker genes *PR10b* and *0s04g10010* [54] in response to 500 nM elf18_*E.coli*_ in mature leaves of 4-week old T_1_ plants derived from *Ubi::EFR::GFP*-1, −7 and −9 and *Ubi::EFR::XA21::GFP* −2, −3 and −4. Only lines expressing EFR::GFP or EFR::XA21::GFP showed induction of *PR10b* and *0s04g10010* expression in response to elf18*_E.coli_*-treatment (Figure 1C and D). The absence of elf18_*E.coli*_-triggered defense gene expression in Kitaake and *Ubi::EFR::GFP-1* plants clearly demonstrates that expression of the EFR-ectodomain is required to confer elf18_*E.coli*_ responsiveness. *PR10b (0s12g36850)* expression was significantly higher in all three lines expressing EFR::XA21::GFP when compared to EFR::GFP expressing lines. This observation is supported by the fact that even the high expression level of EFR::GFP in line 7 does not lead to induction of *PR10b* expression to the same level as in *Ubi::EFR::XA21::GFP* plants (Figure 1B and C).

**Figure 1:**
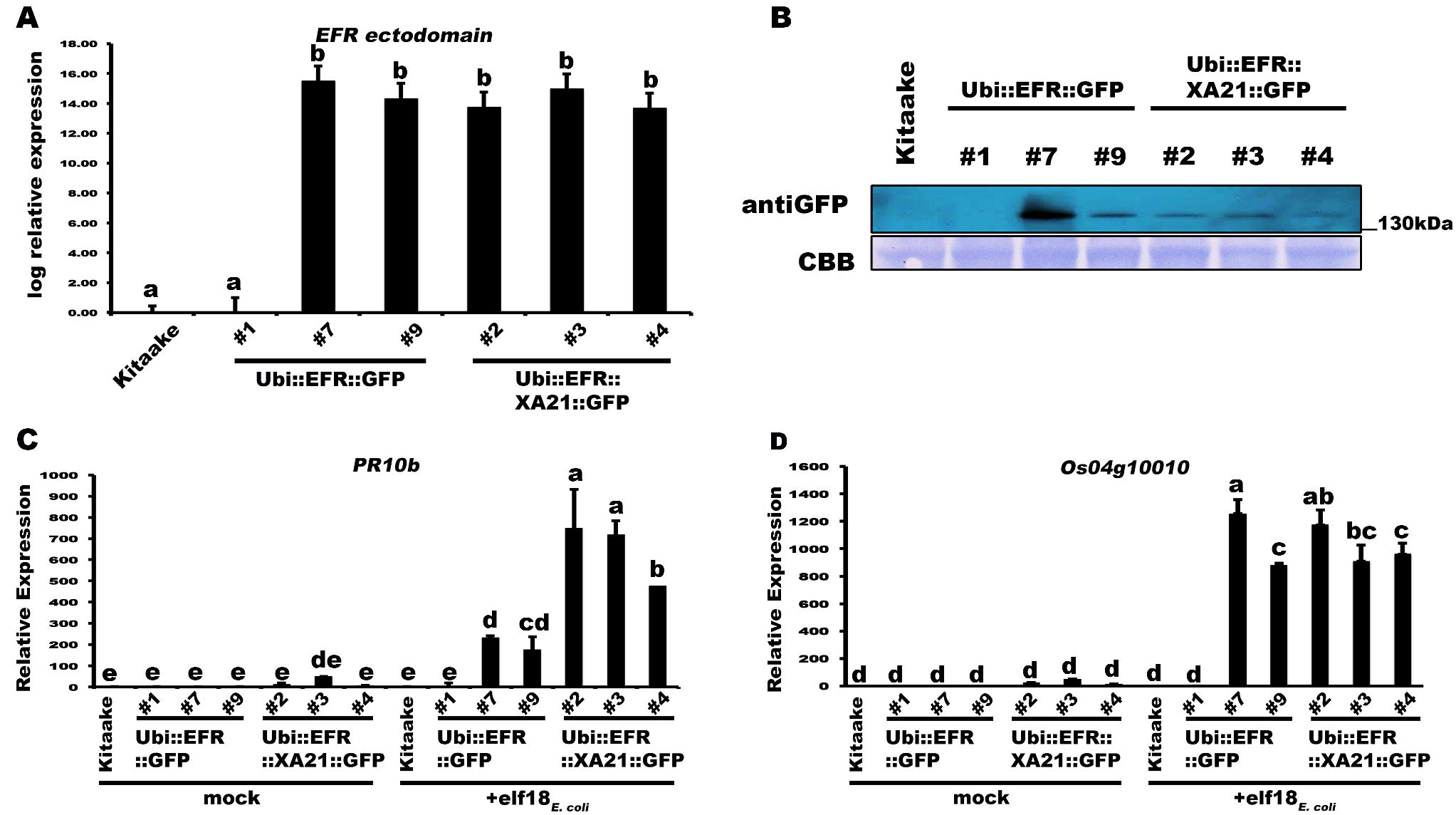
Transgenic expression of EFR and EFR::XA21 in rice leads to elf18_*E.coli*_ responsiveness. (A) Relative expression level of *EFR* and *EFR::XA21* in three independent PCR positive transgenic lines for each immune receptor. Expression was measured by qRT-PCR using primers annealing to the *EFR* ectodomain. Bars depict average expression level relative to actin expression ± SE of three technical replicates. This experiment was repeated at least three times with similar results. (B) Protein level of EFR and EFR::XA21 using an anti-GFP antibody detecting the C-terminal GFP fusion protein. Upper panel anti-GFP western blot, lower panel CBB stain of membrane as loading control. See Supplementary Figure 4 for full western blot. Defense gene expression of *PR10b* (C) and *Os04g10010* (D) in response to elf18_*E.coli*_ (500 nM) in mature leaves of the three independent *Ubi::EFR::GFP* and *Ubi::EFR::XA21::GFP* lines. Expression levels were measured by qRT-PCR and normalized to actin reference gene expression. Data shown is normalized to the Kitaake mock treated (2 hour) sample. Bars depict average expression level ± SE of three technical replicates. This experiment was repeated twice with similar results.

Based on these initial observations, we focused on one line per construct, *Ubi::EFR::GFP-9* and *Ubi::EFR::XA21::GFP-4*, for our next set of experiments (Figure 2 and 3) with the aim of assessing the signaling capacity of both receptor proteins. These and all other transgenic lines used in this study arose from a single T-DNA insertion event as shown by segregation analysis of individual T1 populations (Supplementary Table 1). Because these lines were still segregating in the T_1_ and T_2_ generation we confirmed the presence of the transgene by PCR and performed experiments on PCR positive individuals only. In addition, transgene expression was confirmed by qRT-PCR and found to be stable over multiple generations.

**Figure 2:**
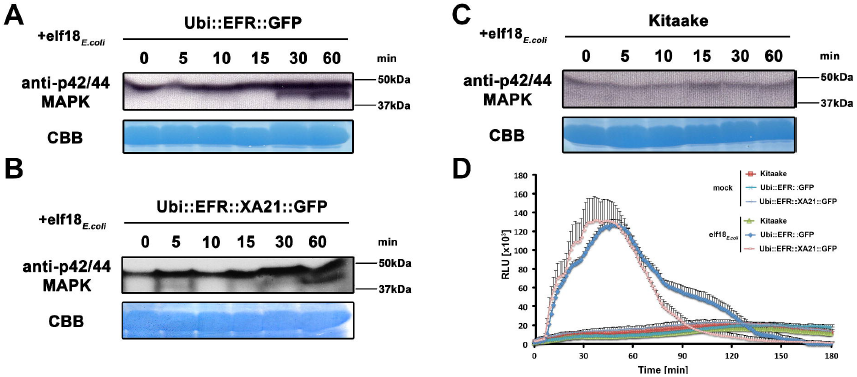
The perception of elf18_*E.coli*_ in EFR and EFR::XA21 rice plants activates several MAP kinases and ROS production. Fully mature leaves of *Ubi::EFR::GFP*-9-11-12 (A), *Ubi::EFR::XA21::GFP-3-6-7* (B) and Kitaake (C), lines were treated with 1 µM *elf18_E.coli_* for the indicated time. Upper panel anti-p42/44 MAP kinase western blot on total protein extracts, lower panel CBB stain of membrane as loading control. (D) Elf18_*E.coli*_-triggered ROS production in fully mature leaves of *Ubi::EFR::GFP*-9-11-12, *Ubi::EFR::XA21::GFP-3-6-7* but not Kitaake. Leaves of the indicated genotypes were treated with water (mock) or *elf18_E.coli_* at a final concentration of 100 nM. Data points depict average relative light production ± SE of at least six biological replicates. These experiment were repeated at least three times with similar results.

**Figure 3:**
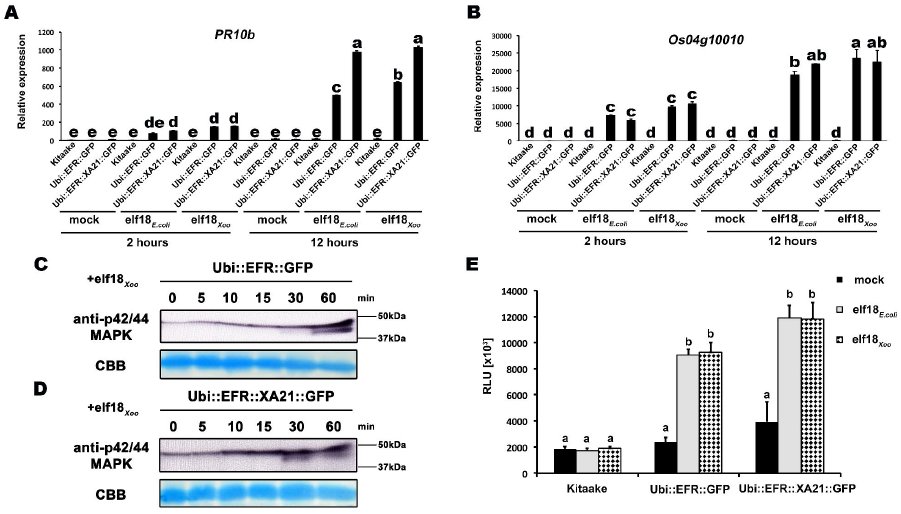
EFR and EFR::XA21 recognize the elf18 sequence derived from *Xoo* EF-Tu. Defense gene expression of *PR10b* (A) and *0s04g10010* (B) in response to elf18_*E.coli*_ or *elf18Xoo* at a concentration of 500 nM in mature leaves of Kitaake, *Ubi::EFR::GFP-9-11-12* and *Ubi::EFR::XA21::GFP-3-6-7* lines. Expression levels were measured by qRT-PCR and normalized to actin reference gene expression. Data shown is normalized to the Kitaake mock treated (2 hour) sample. Bars depict average expression level ± SE of three technical replicates. Fully mature leaves of *Ubi::EFR::GFP-9-11-12* (C) and *Ubi::EFR::XA21::GFP-3-6-7* (D) lines were treated with 1 µM *elf18_E.coli_* for the indicated time. Upper panel anti-p42/44 MAP kinase western blot on total protein extracts, lower panel CBB stain of membrane as loading control. (E) Total ROS production over 3 hours in response to *elf18_E.coli_* or elf18*_Xoo_* at a concentration of 100 nM in mature leaves of Kitaake, *Ubi::EFR::GFP-9*-11-12 and *Ubi::EFR::XA21::GFP-3-6-7* lines. Bars depict average relative light production ± SE of at least six biological replicates. Statistical analysis was performed using the Tukey-Kramer HSD test. Different letters indicate significant differences (p < 0.05). These experiments were repeated at least twice with similar results.

In plants, the kinase domains of several PRRs, including EFR in Arabidopsis, induce MAP kinase activation within minutes of ligand perception [2]. The activation of MAP kinases in Arabidopsis is often measured by an increase of the doubly phosphorylated isoforms detected by an anti-phospho antibody recognizing the two highly conserved activation loop phosphorylation sites pTXpY (where pT or pY represents a phosphorylated threonine or tyrosine, respectively, and x any amino acid) [41]. Recently, this assay has also been established for rice and it was shown that MAP kinases are activated upon treatment with chitin, flg22_*Pta*_, and other MAMPs [61–64]. Before testing the effect of elf18_*E.coli*_ on MAP kinase activation, we first attempted to reproduce MAP kinase activation by flg22*_Pta_* and chitin in mature leaves of 4-week-old Kitaake rice plants treated with 1 μM flg22_*Pta*_ or 50 μg/ml chitin for 0, 5, 10, 15, 30 and 60 minutes. Using the anti-phospho p44/42-antibody, we detected an increase of two distinct bands of an approximate molecular weight of ~47 kDa and 40 kDa after treatment for at least 15 minutes to 30 minutes, respectively (Supplementary Figure 5). The higher molecular weight MAP kinase band was already visible without treatment at 0 minutes, which is in agreement with previous reports [61–64].

Next we tested if elf18_*E.coli*_ treatment of rice leaves expressing EFR::GFP or EFR::XA21 ::GFP would lead to activation of MAP kinases using the same phosphosite-specific antibody. We treated mature leaves of 4-week-old Kitaake, *Ubi::EFR::GFP-9* and *Ubi::EFR::XA21::GFP-4* rice plants with 1 μM elf18_*E.coli*_ for 0, 5, 10, 15, 30 and 60 minutes. We observed a similar activation pattern as observed for chitin and flg22 treatment for both MAP kinases in plants expressing EFR::GFP or EFR::XA21::GFP but not in the Kitaake wild-type plants (Figure 2). The lack of an endogenous receptor for elf18_*E.coli*_ makes Kitaake rice plants insensitive to this elicitor. These observations indicate that the kinase domains of EFR and of XA21 are able to activate MAP kinases in rice in a temporal and ligand-dependent manner.

Similarly to MAP kinase activation, the production of ROS mediated by plasma membrane-localized NADPH oxidases is a frequently utilized measurement to assess PTI signaling [2]. We therefore tested if the expression of EFR and EFR::XA21 in fully mature rice leaves leads to ROS production after treatment with 100 nM elf18_*E.coli*_. We found that only rice plants expressing EFR::GFP or EFR::XA21::GFP triggered ligand-dependent ROS production whereas Kitaake control plants did not respond to elf18_*E.coli*_ treatment (Figure 2D, Supplementary Figure 6). The ROS production peaked around 45 minutes and lasted for about 150 minutes with both lines producing the same amount of total ROS (Figure 2D, Supplementary Figure 6). Using the ROS production assay, we determined the EC50 value of EFR::GFP and EFR::XA21::GFP expressing rice plants. The expression of either receptors confers a highly sensitive chemoperception system for elf18_*E.coli*_ in rice with an EC_50_ value of ~300 pM (Table 1, Supplementary Figure 7), which is very similar to the sensitivity observed in Arabidopsis [27].

**Table 1.** Expression of EFR::GFP and EFR::XA21::GFP in rice confers a high sensitivity to elf18_*E.coli*_ and elf18_*Xoo*_. Table of EC50 values for elf1*8_E.coli_* and elf18_*Xoo*_ in mature leaves of *Ubi::EFR::GFP*-9-11-12 and *Ubi::EFR::XA21::GFP-3-6-7* lines derived from ROS production dose response curves shown in Supplemental Figure 6.

| Genotype | EC50 value |  |
| --- | --- | --- |
|  | *elf18<sub>E.coli</sub>* | *elf18<sub>Xoo</sub>* |
| *Ubi::EFR::GFP* | ~300 pM | ~300 pM |
| *Ubi::EFR::XA21::GFP* | ~300 pM | ~400 pM |

### Rice plants expressing EFR or EFR::XA21 recognize elf18_*Xoo*_ from *Xoo*

Many bacterial species carry two highly similar copies of the *tuf* gene that encodes EF-Tu. In the case of *Xoo*, three full genome sequences are publicly available (PXO99A, KACC10331 and MAFF311018) [65–67]. We investigated the coding sequence for both *tuf* genes, PXO_04524 and PXO_04538, in the *Xoo* PXO99A genome, and for non-synonymous mutations that lead to changes in the 18 N-terminal amino acids (elf18), when compared to the elf18*_E.coli_* amino acid sequence. The elf18*_Xoo_* sequence carried two substitutions when compared to elf18*_E.coli_* (S_2_ –> A_2_ and E_4_ –> A_4_) in both gene copies in all 3 genomes. In addition to the publically available sequences, we analyzed the first ~700 bp of both EF-Tu genes in 20 *Xoo* isolates from our laboratory collection by standard Sanger sequencing (Supplementary Table 2). The first 230 N-terminal amino acids of both EF-Tu proteins were 100% conserved in all 23 *Xoo* isolates giving rise to a single EF-Tu*_Xoo_* sequence as shown in Supplementary Figure 8 [10]. We next tested if elf18*_Xoo_* would be recognized by the EFR ectodomain. We measured defense gene expression in mature leaves of 4-week-old rice plants after treatment with 500 nM elf18*_Xoo_* or elf18_*E.coli*_ for 2 and 12 hours. *PR10b* and *0s04g10010* were up-regulated only in lines expressing EFR::GFP and EFR::XA21::GFP but not the Kitaake control (Figure 3A and B). Both elf18*_Xoo_* and elf18_*E.coli*_ peptides induced a similar defense gene expression levels at all time points and in both lines (Figure 3A and B). As observed previously (Figure 1), the *Ubi::EFR::XA21::GFP* plants induces a higher *PR10b* expression at 12 hours when compared to *Ubi::EFR::GFP* plants. The recognition of elf18*_Xoo_* in mature leaves of 4-week-old rice plants expressing EFR::GFP and EFR::XA21::GFP also induced MAP kinase activation (Figure 3 C and D) and ligand dependent ROS production (Figure 3 E). The sensitivity of EFR and EFR::XA21::GFP plants towards elf18*_Xoo_* was nearly identical with the one observed for elf18_*E.coli*_ with an EC_50_ value of ~300 pM and ~400 pM, respectively (Table 1 and Supplementary Figure 7).

These results suggest that the EFR ectodomain may be able to sense EF-Tu from *Xoo* during the infection process. We therefore tested if EF-Tu*_Xoo_* is present in cell free *Xoo* supernatants. We detected full-length EF-Tu*_Xoo_* in cell free supernatants using a commercially available antibody and by mass-spectrometry analysis (Supplementary Figure 9). This observation is consistent with a recent report that identified EF-Tu*_Xoo_* in the cell-free xylem sap of rice plants infected with *Xoo* [68]. Therefore, we hypothesize that EF-Tu from *Xoo* is readily available for detection by EFR and EFR::XA21 during the infection process.

### Transgenic expression of EFR in rice does not negatively affect growth and yield

In some instances, transgenic expression of defense related genes has a negative impact on the plant’s growth and yield. To test whether the transgenic expression of EFR or EFR::XA21 has a negative impact on rice growth and yield, we grew wild type Kitaake plants next to *Ubi::EFR::GFP* and *Ubi::EFR::XA21::GFP* lines until maturity and measured total dry biomass and yield. Supplementary Figure 10 shows that two independent transgenic rice lines expressing EFR *(Ubi::EFR::GFP-7-8-8* and Ubi::EFR::GFP-9-4-3-13) do not differ in total biomass or yield compared to the wild type parent. In contrast, EFR::XA21::GFP expressing plants *(Ubi::EFR::XA21::GFP-3-8-7-20* and *Ubi::EFR::XA21::GFP-4-5-4)* do suffer from growth defects such as necrosis of older leaves and stunting starting at the 5-week stage under our greenhouse conditions. Although the overall biomass and yield of EFR::XA21::GFP plants was reduced at maturity (Supplementary Figure 10), the *Ubi::EFR::XA21::GFP* plants did not show any macroscopic necrotic lesions or early senescence until the 5-week-old stage.

### Transgenic expression of EFR::XA21 in rice does not alter steady-state defense gene expression in 4-week-old plants

We investigated by RNA sequencing if *Ubi::EFR::XA21::GFP* plants exhibited stress-related symptoms even before the onset of necrotic lesions. The transcriptomic profile of mature leaves of 4-week-old *Ubi::EFR::XA21::GFP-3-4* and Kitaake soil grown plants were compared to determine if stress-related genes were differentially regulated in *Ubi::EFR::XA21::GFP* plants. First, we investigated if gene expression patterns of the different genotypes are distinct. Pearson correlation coefficients show that replicates from Kitaake and *Ubi::EFR::XA21::GFP* cluster together in pairwise analysis (Supplementary Figure 11). We identified 131 genes, which were differentially expressed between Kitaake and *Ubi::EFR::XA21::GFP* plants with a median log fold change of 2.94, using a false discovery rate of ≤ 0.05 and an absolute log fold-change ≥ 2.00 (Supplementary Table 3). This differential gene expression list includes 115 up-regulated and 26 down-regulated genes. The defense marker gene *Os04g10010* was not included in the set of up-regulated genes in *Ubi::EFR::XA21::GFP* plants in the absence of the ligand treatment consistent with our previous observations (Figure 1 and 3). In contrast, *PR10b* expression was up-regulated by 2.17-fold, which is just above our 2.00-fold log fold-change cut-off (Supplementary Table 3). We observed a similar slight up-regulation of *PR10b* in *Ubi::EFR::XA21::GFP* plants in the absence of ligand treatment in other experiments, however this slight up-regulation of *PR10b* in *Ubi::EFR::XA21::GFP* plants was not statistically significant (Figure 1A and 3B). Similarly, none of these defense marker genes was differentially expressed in the absence of the ligand in any experiments performed with *Ubi::EFR::GFP* rice plants (Figure 1 and 3).

Gene ontology (GO) term analyses using the up-regulated gene set of *Ubi::EFR::XA21::GFP* plants showed no significant enrichment for any GO terms (Supplementary Figure 12A). GO term analysis using the down-regulated gene set of *Ubi::EFR::XA21::GFP* plants showed significant enrichment (6 out of 26) for the GO term ‘oxidoreductase activity’ (p = 0.023, FDR = 0.042) (Supplementary Figure 12B). The whole transcriptome analysis of *Ubi::EFR::XA21::GFP* plants at the 4-week-old stage in the absence of the ligand indicated that *Ubi::EFR::XA21::GFP* plants do not overexpress stress-related genes at this plant stage. Indeed the transcriptomes of *Ubi::EFR::XA21::GFP* plants and the Kitaake control plants are nearly identical with an overall correlation coefficient of R > 0.99.

### Transgenic rice plants expressing the EFR receptor are more resistant to two weakly virulent *Xoo* isolates

The expression of EFR and EFR::XA21 enables rice to sense and respond to elf18*_Xoo_* at sub-nanolmolar concentrations (Figure 3, Table 1). Moreover, EF-Tu*_Xoo_* is most likely readily available for EFR recognition during the infection process (Supplementary Figure 9) [69]. To determine whether the transgenic expression of EFR or EFR::XA21 in rice confers enhanced resistance to *Xoo*, we inoculated *Ubi::EFR::GFP* and *Ubi::EFR::XA21::GFP* transgenic plants and compared the length of disease lesions with that of Kitaake plants. EFR expression did not confer robust resistance to the fully virulent PXO99A isolate. We tested three independent EFR lines: **3**–6 (T1), **7**–8–8 (T2) and **9**–11–2 (T2)/**9**–4–3–13 (T3), referred to as lines 3, 7 and 9, respectively, as detailed in Table 2. In 5 out of 8 infections of *Ubi::EFR::GFP* plants with PXO99A, we observed a moderate, but statistically significant, reduction in lesion length as shown in Figure 4A. *In planta* bacterial growth curve analysis revealed no statistical difference in PXO99A populations between EFR and Kitaake lines in three independent experiments (Supplementary Figure 13). When we attempted to perform similar inoculation assays with *Ubi::EFR::XA21::GFP* lines, we had difficulties maintaining healthy plants throughout the course of inoculation starting at the 6-week-old stage (Supplementary Figure 10). However, in two experiments we did obtain healthy plants and carried out the inoculation assays in full. In these assays (see experiment number II and number VI in Table 2) *Ubi::EFR::XA21::GFP* plants were as susceptible to *Xoo* PXO99A infection as Kitaake control plants. These results indicate that despite the ability of the EFR and EFR::XA21 chimeric receptors to detect EF-Tu*_Xoo_* in the detached leaf assay (Figure 3), this recognition does not confer robust resistance to *Xoo* PXO99A in whole rice plants.

**Figure 4:**
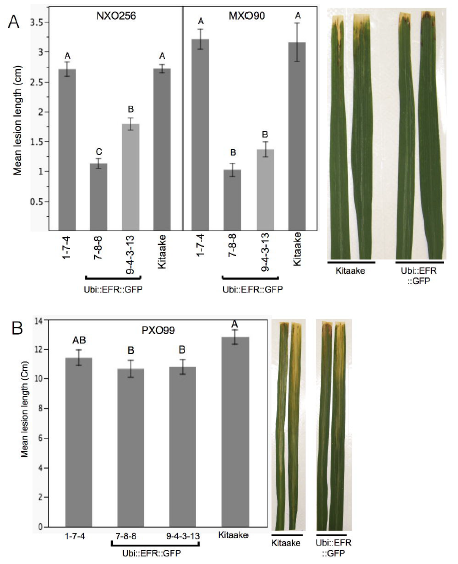
Rice lines expressing the EFR receptor are quantitatively more resistant to weakly virulent *Xoo* isolates. (A) Two independent *Ubi::EFR::GFP*-7 and −9 lines expressing EFR and Kitaake control were inoculated with *Xoo* PXO99A. (B) Two independent *Ubi::EFR::GFP-7* and -9 lines expressing EFR, a null transgene *Ubi::EFR::GFP-1* control and wild-type Kitaake were inoculated with two weakly virulent isolates (NXO256 and MXO90), (see experiment V in infection summary Table 2). Plants were infected using the leaf-clipping assay at the 5-to 6-week-old stage. Lesions were measured at 14 dpi. On the right hand side of each panel two representative inoculated leaves at the time of lesion scoring for either Kitaake or *Ubi::EFR::GFP* transgenic line. Statistical analysis was performed using the Tukey-Kramer HSD test. Different letters indicate significant differences within each Isolate. These experiments were repeated at least twice with similar results.

Because the leaf-clipping infection assay might mask subtle differences in resistance, we sought to use less aggressive infection assays. We took two approaches to address this issue. In our first approach we inoculated plants with a lower dose of the fully virulent PXO99A isolate. In our second approach we inoculated with weakly virulent *Xoo* isolates. Inoculation with a lower concentration of the PXO99A isolate (10^6^ CFU/mL instead of 10^8^ CFU/mL) did not result in statistically significant differences in disease lesion length between wild type and EFR transgenic plants (Supplementary Table 4). We next screened 10 different *Xoo* isolates from our lab collection for their level of virulence on Kitaake plants. We identified three isolates that were significantly less virulent than PXO99A and other fully virulent isolates (Supplementary Table 4). Two of these weakly virulent isolates (NXO256 and MXO90), were significantly less virulent on rice lines expressing EFR when compared with wild-type Kitaake plants (Figure 4 B, Table 2). *Ubi::EFR::GFP* lines (7 and 9) were statistically significantly more resistant to isolate MXO90 (shorter lesions) in 8 out of 8 inoculations and more resistant to isolate NXO256 in 5 out of 6 inoculations (Table 2). These moderately enhanced resistance phenotypes were caused by the expression of EFR in the *Ubi::EFR::GFP* lines. The rice line *Ubi::EFR::GFP-* 1, which does not express EFR, is not responsive to elf18_*E.coli*_ (Figure 1) and is not resistant to *Xoo* (Figure 4, Table 2 experiment V). The moderate resistance conferred by the *Ubi::EFR::GFP* lines, as measured by lesion lengths, was further supported by *in planta* growth curves (Supplementary Figure 13). These results indicate that the recognition of elf18*_Xoo_* by EFR leads to a reduction of bacterial populations of these two weakly virulent isolates. When we tested the *Ubi::EFR::XA21::GFP* lines with these two isolates (see experiment number II and number VI in Table 2), the lesion lengths were between those obtained with Kitaake and those obtained with *Ubi::EFR::GFP* transgenic lines. In the case of the *Xoo* isolate NXO256, no significant differences in lesion length could be observed on *Ubi::EFR::GFP* transgenic lines. For the *Xoo* isolate MXO90, statistically significant differences between Kitaake and *Ubi::EFR::GFP* transgenic lines were observed 1 out 2 experiments (Table 2).

**Table 2.**
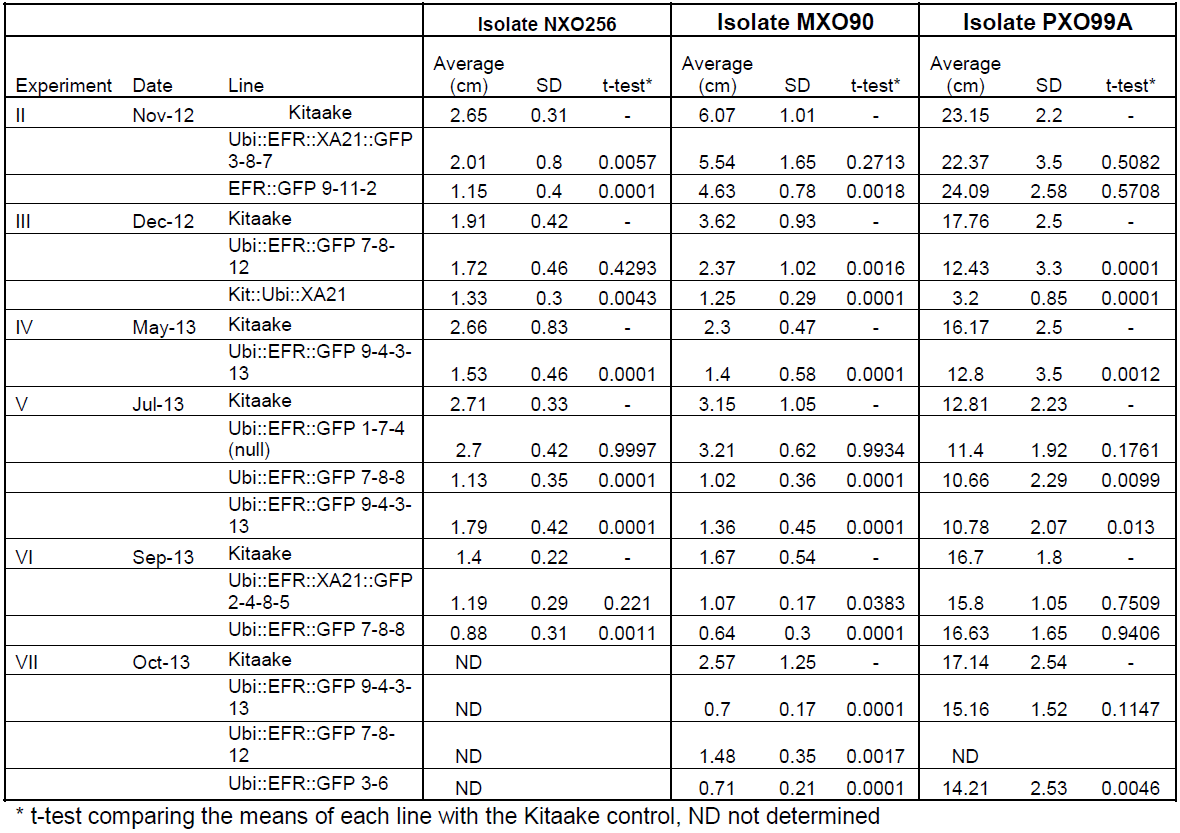
Summary of *X*. oryzae pv. oryzae (*Xoo*) inoculation experiments. Detailed description of all six inoculation experiments comparing Kitaake control plants compared to *Ubi::EFR::GFP* and *Ubi::EFR::XA21::GFP* transgenic rice lines following Xoo inoculation using isolate PXO99A (fully virulent) and isolates NXO256 and MXO90 (weakly virulent).

|  |  |  | Isolate NXO256 |  |  | Isolate MXO90 |  |  | Isolate PXO99A |  |  |
| --- | --- | --- | --- | --- | --- | --- | --- | --- | --- | --- | --- |
| Experiment | Date | Line | Average (cm) | SD | t-test* | Average (cm) | SD | t-test* | Average (cm) | SD | t-test* |
| II | Nov-12 | Kitaake | 2.65 | 0.31 | - | 6.07 | 1.01 | - | 23.15 | 2.2 | - |
|  |  | Ubi::EFR::XA21::GFP 3-8-7 | 2.01 | 0.8 | 0.0057 | 5.54 | 1.65 | 0.2713 | 22.37 | 3.5 | 0.5082 |
|  |  | EFR::GFP 9-11-2 | 1.15 | 0.4 | 0.0001 | 4.63 | 0.78 | 0.0018 | 24.09 | 2.58 | 0.5708 |
| III | Dec-12 | Kitaake | 1.91 | 0.42 | - | 3.62 | 0.93 | - | 17.76 | 2.5 | - |
|  |  | Ubi::EFR::GFP 7-8-12 | 1.72 | 0.46 | 0.4293 | 2.37 | 1.02 | 0.0016 | 12.43 | 3.3 | 0.0001 |
|  |  | Kit::Ubi::XA21 | 1.33 | 0.3 | 0.0043 | 1.25 | 0.29 | 0.0001 | 3.2 | 0.85 | 0.0001 |
| IV | May-13 | Kitaake | 2.66 | 0.83 | - | 2.3 | 0.47 | - | 16.17 | 2.5 | - |
|  |  | Ubi::EFR::GFP 9-4-3-13 | 1.53 | 0.46 | 0.0001 | 1.4 | 0.58 | 0.0001 | 12.8 | 3.5 | 0.0012 |
| V | Jul-13 | Kitaake | 2.71 | 0.33 | - | 3.15 | 1.05 | - | 12.81 | 2.23 | - |
|  |  | Ubi::EFR::GFP 1-7-4 (null) | 2.7 | 0.42 | 0.9997 | 3.21 | 0.62 | 0.9934 | 11.4 | 1.92 | 0.1761 |
|  |  | Ubi::EFR::GFP 7-8-8 | 1.13 | 0.35 | 0.0001 | 1.02 | 0.36 | 0.0001 | 10.66 | 2.29 | 0.0099 |
|  |  | Ubi::EFR::GFP 9-4-3-13 | 1.79 | 0.42 | 0.0001 | 1.36 | 0.45 | 0.0001 | 10.78 | 2.07 | 0.013 |
| VI | Sep-13 | Kitaake | 1.4 | 0.22 | - | 1.67 | 0.54 | - | 16.7 | 1.8 | - |
|  |  | Ubi::EFR::XA21::GFP 2-4-8-5 | 1.19 | 0.29 | 0.221 | 1.07 | 0.17 | 0.0383 | 15.8 | 1.05 | 0.7509 |
|  |  | Ubi::EFR::GFP 7-8-8 | 0.88 | 0.31 | 0.0011 | 0.64 | 0.3 | 0.0001 | 16.63 | 1.65 | 0.9406 |
| VII | Oct-13 | Kitaake | ND |  |  | 2.57 | 1.25 | - | 17.14 | 2.54 | - |
|  |  | Ubi::EFR::GFP 9-4-3-13 | ND |  |  | 0.7 | 0.17 | 0.0001 | 15.16 | 1.52 | 0.1147 |
|  |  | Ubi::EFR::GFP 7-8-12 | ND |  |  | 1.48 | 0.35 | 0.0017 | ND |  |  |
|  |  | Ubi::EFR::GFP 3-6 | ND |  |  | 0.71 | 0.21 | 0.0001 | 14.21 | 2.53 | 0.0046 |

\* t-test comparing the means of each line with the Kitaake control, ND not determined

In summary, while the expression of EFR and EFR::XA21 does not confer robust resistance to fully virulent *Xoo* isolate PXO99A, expression of EFR provides quantitatively enhanced resistance to two weakly virulent *Xoo* isolates.

### EFR interacts with OsSERK2 and XB24, but not with XB3 and XB15

Transgenic expression of the Brass/caceae-s pecific PRR EFR in rice confers both sensitivity to elf18*_Xoo_*/elf18_*E.coli*_ and quantitatively enhanced resistance against two weakly virulent *Xoo* isolates (Figures 1–4, Table 1 and 2). We therefore hypothesized that EFR in rice engages at least a subset of XA21-signaling network components [70]. To test this hypothesis, we investigated the interaction of EFR with four major XA21 interaction partners [53,54,56,59]. We performed targeted yeast two-hybrid experiments between the EFR intracellular domain (ID) (674-1032aa) and OsSERK2 ID (260-628aa), XB3 full-length (FL) (1-450aa), XB15 FL (1-639aa) and XB24 FL (1-198aa) [53,54,56,59].

We found that the XA21 ID (668-1025aa) interacted with all four proteins (Supplementary Figure 14) as previously reported [53,54,56,59]. Next, we tested the interaction of EFR ID with the same four proteins. In these experiments, the EFR ID interacted with XB24 but not XB3, XB15 or OsSERK2 ID (Figure 5A). The expression of all fusion proteins in yeast was confirmed by western blot analysis (Supplementary Figure 15). The interaction of XA21 with XB24 is dependent on the catalytic activity of the XA21 kinase [53]. We tested if this is also the case for the EFR/XB24 interaction by yeast-two hybrid analysis between catalytically inactive EFR(D848N) ID and XB24. For this purpose, we mutated the conserved aspartate at position 848 in EFR, which was previously shown to be required for catalytic activity [41], to an asparagine. In the yeast two-hybrid system, EFR(D848N) was still able to directly interact with XB24 (Figure 5A). This suggests that the interaction between EFR and XB24 is independent of the kinase catalytic activity of EFR. Next, we aimed to confirm the interaction between XB24 and EFR *in planta*. In the absence of a suitable antibody for XB24, we decided to test this interaction in *N. benthamiana* after co-expression of tagged versions of each protein using *Agrobacterium tumefaciens-mediated* transient expression. As shown in Figure 5B, XB24::FLAG specifically co-immunoprecipitated with EFR::GFP. The association was unaltered by treatment with 100 nM elf18*_E.coli_* for 10 minutes.

**Figure 5:**
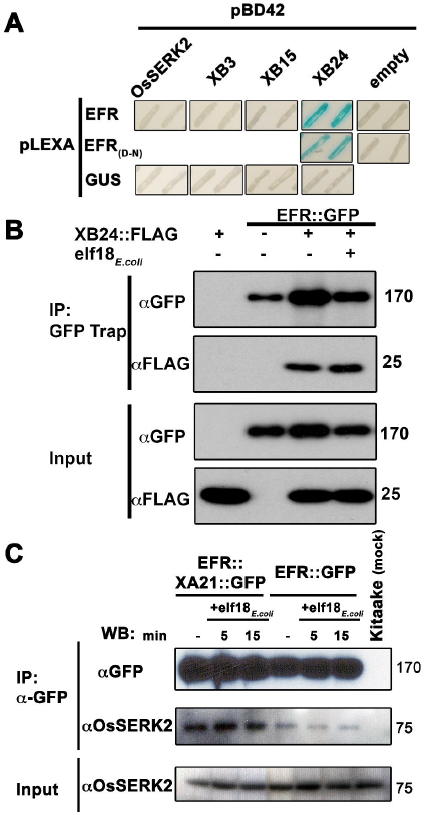
EFR interacts with the XA21-signaling components XB24 and OsSERK2. (A) Yeast two-hybrid assay between EFR (674-1032aa) intracellular domain (ID) and OsSERK2 ID (260-628aa), XB3 full-length (FL) (1-450aa), XB15 FL (1-639aa) and XB24 FL (1-198aa). EFR(D-N) indicates a mutation of the catalytic aspartate (D) 848aa to asparagine (N). The blue color indicates nuclear interaction between the two co-expressed proteins. Expression of all fusion proteins was confirmed by western blot analysis as shown in Supplementary Figure 15. (B) *In planta* interaction of EFR::GFP and XB24::FLAG in *N. benthamiana.* EFR::GFP and XB24::FLAG were transiently co-expressed in *N. benthamiana* leaves. Leaves were mock treated with water (-) or with 100 nM elf18_*E.coli*_ for 10 min. Immuno-complexes were precipitated (IP) with GFP-Trap agarose beads. Proteins were separated and detected by SDS-PAGE electrophoresis followed by western blot analysis with HRP-conjugated anti-GFP or anti-FLAG. (C) EFR::GFP or EFR::XA21::GFP form constitutive ligand-independent complexes with OsSERK2 *in vivo* without quantitative changes of the interaction within 15 min of elf1*8_E.coli_* treatment. Immuno-complexes were precipitated from leaf material of *Ubi::EFR::GFP-9*-11-12 and *Ubi::EFR::XA21::GFP-3-6-7* expressing rice plants treated with 1 μM elf18_*E.coli*_ for the indicated time using GFP-trap beads. Kitaake rice leaves were used as negative control. Components of the immuno-precipitated complexes were separated by SDS-PAGE gel followed by immuno-detection with anti-GFP (for EFR::GFP and EFR::XA21::GFP) and anti-OsSERK2 (for OsSERK2). EFR::GFP and EFR::XA21::GFP gives rise to a signal at about 175 kDa. OsSERK2 (~70 kDa) was co-immunoprecipitated with EFR::GFP and EFR::XA21::GFP in the absence of elf18 treatment. The lower panel shows equal amounts of OsSERK2 in both total protein fractions before immunoprecipitation. This experiment was repeated twice with similar results.

The absence of a direct interaction between the EFR ID and OsSERK2 ID in the yeast two-hybrid system is surprising because orthologous SERK family members in rice and Arabidopsis interact with XA21 or EFR *in vivo and* are required for XA21-and EFR-mediated immune responses [37,40–42,54]. We therefore hypothesized that we may be able to detect the interaction *in planta*. For this purpose, we used our recently developed specific anti-OsSERK2 antibody [54], to test for the interaction between EFR::GFP and EFR::XA21::GFP in leaf strips of 4-week-old plants after treatment with 1 µM *elf18_E.coli_* or elf18*_Xoo_* for 0, 5 and 15 minutes. We choose these time points based on previous interaction data reported for EFR-AtSERK3/BAK1(At4g33430) at 5 minutes [38,41,42] and on *in vivo* data demonstrating initial MAP kinase activation within 15 minutes of elf18*_E.coli_* treatment (Figure 2). In immunoprecipitation experiments with anti-GFP agarose, we detected proteins at the expected size of 175kDa for full-length EFR::GFP and EFR::XA21::GFP using anti-GFP antibody only in transgenic plants and not in Kitaake control plants (Figure 5B). Next, we tested for the presence of OsSERK2 in the anti-GFP immunoprecipitates using anti-OsSERK2 antibody. OsSERK2 was readily detectable in all immunoprecpitates from EFR::GFP or EFR::XA21::GFP expressing plants even in the absence of elf18 treatment but not in immunoprecipitates from Kitaake control plants (Figure 5B). No increase in co-immunoprecipitated OsSERK2 could be observed within 15 minutes of elf18_*E.coli*_ treatment (Figure 5B). This is consistent with our previous observation that XA21 and OsSERK2 form constitutive heteromeric complexes in the same plant tissue [54]. In contrast, the interaction between EFR and AtSERK3/BAK1 is clearly ligand-induced in Arabidopsis and after transient coexpression in *N. benthamiana* [41,42]. These interaction studies indicate that EFR in rice utilizes at least a subset of the XA21-signaling components.

### OsSERK2 and XB24 regulate EFR signaling in rice

Based on the interaction of EFR with OsSERK2 and XB24, we tested if OsSERK2 and XB24 are also involved in EFR-signaling in rice by assessing double transgenic lines of *Ubi::EFR::GFP* with altered expression of *OsSERK2* and *XB24.* Because OsSERK2 directly interacts with EFR, we tested if OsSERK2 is also a positive regulator of EFR signaling in rice. We crossed *Ubi::EFR::GFP-9-2* expressing lines with previously characterized *OsSERK2RNAi* silencing lines [54]. In the F2 generation, we isolated double transgenic lines from two independent F1 plants (67 and 71) by PCR using primers specific for each transgene and confirmed stable expression of the EFR transgene by qRT-PCR (Supplementary Figure 16A). We compared elf18_*E.coli*_-induced defense gene expression in plants expressing EFR::GFP and silenced for *OsSERK2 (Ubi::EFR::GFP* x *OsSERK2RNAi*) with plants expressing only EFR::GFP (*Ubi::EFR::GFP*). We treated leaf strips of 4-week-old plants from both genotypes with water or 500 nM elf18_*E.coli*_ for 2 and 12 hours. As shown in Figure 6A and B, elf18*_E.coli_*-induced *PR10b* and *0s04g10010* expression was significantly reduced in both independent double transgenic lines *Ubi::EFR::GFP* x *OsSERK2RNAi-67* and −71 expressing EFR::GFP and silenced for *OsSERK2* at both time points when compared with EFR::GFP expressing controls. Similarly, the ROS production triggered by 100 nM elf18_*E. coli*_ application was significantly reduced and delayed in EFR::GFP expressing lines silenced for *OsSERK2* (Figure 6C and D). These results demonstrate that silencing of *OsSERK2* interferes with elf18_*E.coli*_-induced EFR signaling in rice. Similar to AtSERK3/BAK1 and AtSERK4/BKK1 in Arabidopsis, OsSERK2 appears to be a positive regulator of EFR signaling in rice orthologous to its role in XA21 signaling [42,54].

**Figure 6:**
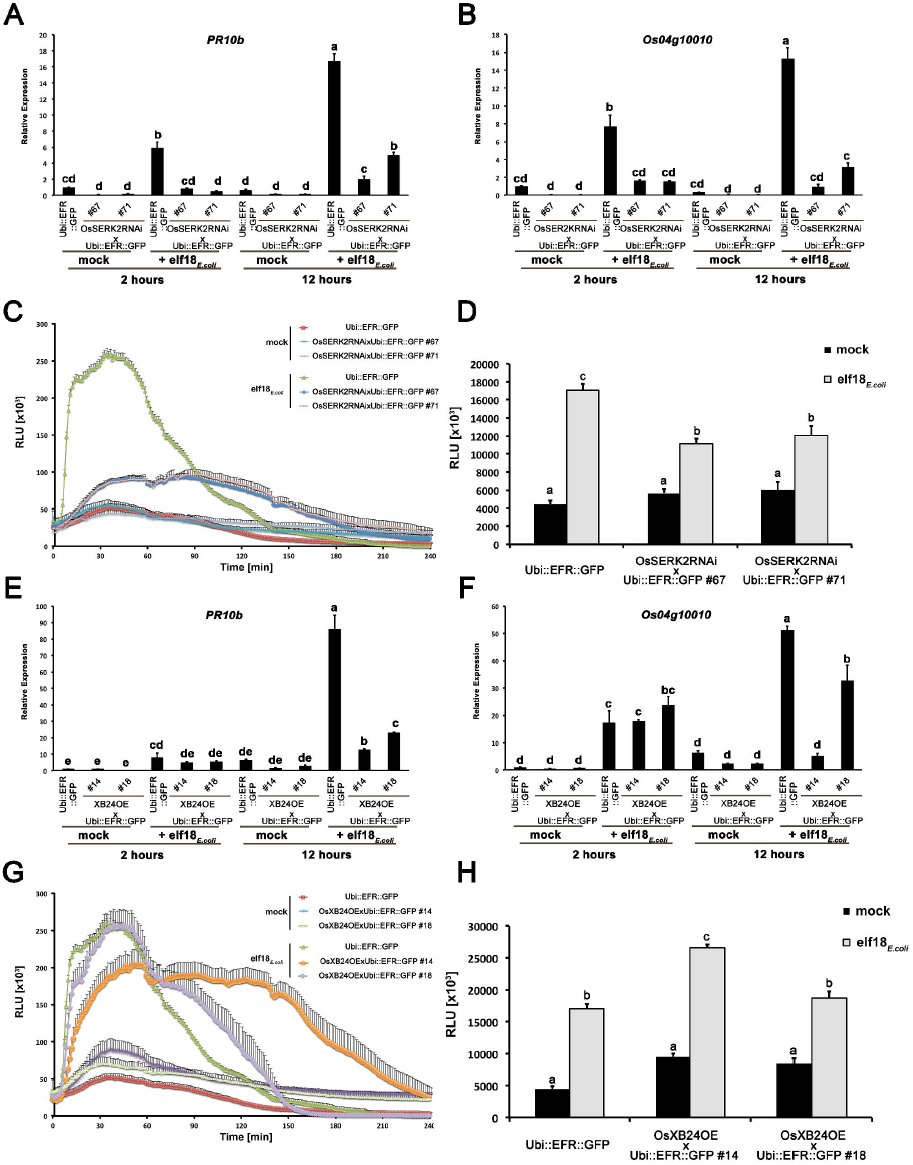
*OsSERK2* and *XB24* are involved in EFR signaling in rice. Defense gene expression of *PR10b* (A) and *0s04g10010* (B) in response to elf18_*E.coli*_ at a concentration of 500 nM in mature leaves of *Ubi::EFR::GFP*-9-11-12 and double transgenic F2 (67 and 71) plants from two independent crosses between *Ubi::EFR::GFP*-9-11 and *OsSERK2RNAi-X-B-4-2*. Expression levels were measured by qRT-PCR and normalized to actin reference gene expression. Data shown is normalized to the Kitaake mock treated (2 hour) sample. Bars depict average expression level ± SE of three technical replicates. (C) and (D) Temporal and total ROS production over 4 hours in response to elf18_*E.coli*_ or elf18*_Xoo_* at a concentration of 100 nM in mature leaves of *Ubi::EFR::GFP*-9-11-12 and double transgenic F2 (67 and 71) plants from two independent crosses between *Ubi::EFR::GFP*-9-11 and *OsSERK2RNAi-X-B-4-2.* Data points or bars depict average relative light production ± SE of at least six biological replicates. Defense gene expression of *PR10b* (E) and *0s04g10010* (F) in response to elf18_*E.coli*_ at a concentration of 500 nM in mature leaves of *Ubi::EFR::GFP*-9-11-12 and double transgenic F2 (14 and 18) plants from two independent crosses between *Ubi::EFR::GFP-9-11* and *XB24* overexpressing (OE) line A109-6-5-1. Expression levels were measured by qRT-PCR and normalized to actin reference gene expression. Data shown is normalized to the Kitaake mock treated (2 hour) sample. Bars depict average expression level ± SE of three technical replicates. (G) and (H) Temporal and total ROS production over 4 hours in response to elf18_*E.coli*_-or elf18*_Xoo_* at a concentration of 100 nM in mature leaves of *Ubi::EFR::GFP*-9-11-12 and double transgenic F2 (14 and 18) plants from two independent crosses between *Ubi::EFR::GFP*-9-11 and *XB24* overexpressing (OE) line A109-6-5-1. Data points or bars depict average relative light production ± SE of at least six biological replicates.

In contrast to OsSERK2, XB24 is a negative regulator of XA21-mediated immunity [53]. To test if XB24 is also involved in EFR signaling in rice, we crossed previously described *XB24* overexpressing lines *(XB24OE A109-6-5-1)* [53] with lines expressing EFR::GFP *(Ubi::EFR::GFP-9-2*). In the F2 generation, we isolated double transgenic lines from two independent crosses (14 and 18) by PCR with primers specific for each transgene and confirmed stable expression of the EFR transgene by qRT-PCR (Supplementary Figure 16B). We assessed the impact of XB24 overexpression on elf18_*E.coli*_-induced EFR-signaling in defense gene expression and ROS assays as describe above. Elf18_*E.coli*_-triggered defense gene expression was significantly reduced at 12 hours post treatment for both marker genes in both lines overexpressing *XB24* in the EFR::GFP background (*Ubi::EFR::GFP x XB24OE-14* and -18) when compared to EFR::GFP expressing controls (Figure 6E and F). Similarly, ROS production was significantly altered with both lines showing an extended ROS production (Figure 6G). This effect was more pronounced in the line *Ubi::EFR::GFP x XB24OE-14* and led to a significant increase in total ROS production in this line only (Figure 6H). This is consistent with the more severe defect of elf18_*E.coli*_-triggered defense gene expression in this line (Figure 6E and F). This suggests that XB24 is involved in EFR signaling in rice when overexpressed, similarly to its negative role in XA21-signaling.

## Discussion

The main aim of this study was to investigate the feasibility of inter-class transfer of dicotolydenous PRRs, such as EFR from Arabidopsis, into the model monocotyledonous species, rice. We aimed to determine if PRRs, when transgenically expressed, from evolutionarily distant plant species can enhance resistance and if they employ the same signaling components as endogenous PRRs. We demonstrate that transgenic expression of EFR in rice confers a high-level of sensitivity to the previously unrecognized elf18*_E.coli_* and elf18*_Xoo_*. Ligand-dependent activation of EFR elicits well-characterized defense responses such as defense gene expression, ROS production and MAP kinase activation (Figures 1–3, Table 1). While expression of EFR does not lead to robust enhanced resistance to fully virulent *Xoo* isolates, it does lead to quantitatively enhanced resistance to two weakly virulent *Xoo* isolates (Figure 4, Table 2). We made similar observations of full defense response activation at sub-nanomolar concentrations of elf18_*E.coli*_ and elf18*_Xoo_* (Table 1) but limited enhanced resistance to *Xoo* for rice plants expressing the chimeric receptor EFR::XA21, which consists of the EFR ectodomain and the intracellular XA21 kinase domain (Figure 1–4, Table 1 and 2). These results indicate that ligand activated XA21 kinase alone is not sufficient to induce the robust resistance response observed with the full-length XA21.

EFR in rice utilizes at least two well-described XA21-signaling components OsSERK2 and XB24, most likely via direct protein interactions (Figure 5 and 6).

### Signaling downstream of EFR and EFR::XA21 in rice

In recent years, tremendous advances have been made in deciphering the signaling events occurring at the PRR level within seconds to minutes of ligand perception [2,15,71]. These advances have been mainly driven by studies in Arabidopsis involving EFR, FLS2 and CERK1 and in rice involving chitin perception by CEBiP [2,15,71]. Most of the recent progress on rice receptor kinase PRRs, including XA21 and XA3, has relied on the characterization of much later phenotypic read-outs such as disease progression, which is recorded over 1 week after inoculation [72,73]. This is mostly caused by the paucity of well-defined ligands for most rice receptor kinase PRRs such as XA21, Pi-d2 (FJ915121.1) and XA3 (DQ426646.1) [74–77]. In Arabidopsis, several well-defined defense read-outs are readily available to assess signaling activation post-ligand treatment including defense marker genes, ROS burst and MAP kinase activation [2]. Very little is known about the immediate signaling activated by peptide ligands in the absence of infectious agents in fully mature rice leaves.

We used EFR and the chimeric receptor EFR::XA21 to probe rice responses using the well-defined peptide ligand elf18*_E.coli_*. Both receptors elicit qualitatively similar defense signaling pathways including the activation of several MAP kinases, ROS production, and up-regulation of two defense maker genes, *PR10b* and *0s04g10010* (Figure 1 to 3, Table 1). Indeed expression of EFR::GFP and EFR::XA21::GFP confers similar, high sensitivity to elf18_*E.coli*_ and elf18*_Xoo_* as was previously reported for EFR in Arabidopsis with an EC_50_ value of ~300 pM (Table 1). While both receptors elicited both marker genes, the kinase domain of XA21 appears to consistently lead to a higher up-regulation of *PR10b* (Figure 1C and 3B). This suggests that the XA21 kinase might be better adapted to defense signaling in rice as compared with the EFR kinase domain, which is derived from a dicotyledonous plant species. This increased signal capacity of the XA21 kinase domain might also be the reason why older *Ubi::EFR::XA21::GFP* plants appear to be necrotic, tend to senesce earlier and accumulate lower biomass at full maturity (Supplementary Figure 10). These phenotypes of *Ubi::EFR::XA21::GFP* plants might be caused by the abundance of EF-Tu in the rhizosphere and phyllosphere. This detection might lead to a stronger, more severe continuous defense activation in *Ubi::EFR::XA21::GFP* plants when compared with *Ubi::EFR::GFP* plants.

The observed severe phenotypic differences between *Ubi::EFR::XA21::GFP* versus *Ubi::EFR::GFP* and Kitaake controls were clearly age-dependent and only observable from the 5-week-old stage onwards. When we attempted to identify underlying signaling pathways that may be activated at the 4-week-old stage, before macroscopic necrosis developed, we only detected 131 differentially expressed genes when comparing *Ubi::EFR::XA21::GFP* with Kitaake control in the absence of ligand treatment (Supplementary Table 3). No specific GO terms were enriched in the up-regulated gene set (Supplementary Figure 12A). However, we identified a significant enrichment for GO terms associated with oxidoreductase activity in genes down-regulated in EFR::XA21 (Supplementary Figure 12B). This suggests that *Ubi::EFR::XA21::GFP* plants might be more susceptible to oxidative stress at older developmental stages. Consistent with these observations, SA signaling appears to be activated in these 5-week-old plants as two SA marker genes are upregulated in EFR::XA21::GFP expressing plants in the absence of any treatment (Supplementary Figure 17). However, the overall transcriptomic comparison between *Ubi::EFR::XA21::GFP* and Kitaake suggests that at 4-week-old stage *Ubi::EFR::XA21::GFP* plants are very similar to wild-type plants and do not overexpress stress related genes. These results indicate that the *Ubi::EFR::XA21::GFP* plants serve as a useful surrogate system to investigate the transcriptional reprogramming induced by ligand activated XA21 kinase in rice leaf tissue.

Our studies to determine if EFR in rice utilizes similar signaling components as XA21 [2,70–72] identified OsSERK2 and XB24 but not XB3 and XB15 as interaction partners of EFR (Figure 5). We therefore focused our further genetic studies on *OsSERK2* and *XB24.* We crossed EFR-expressing rice lines to *OsSERK2*-silenced lines and *XB24*-overexpressing lines. We found that OsSERK2 is a positive regulator of EFR-mediated defense signaling in rice, similar to its role in the Xa21-mediated immune response [54]. EFR lines silenced for *OsSerk2* are significantly impaired in elf18-induced defense gene expression and ROS production (Figure 6A to 6D). In Arabidopsis, EFR requires several SERK proteins for its function including SERK3/BAK1 and SERK4/BKK1 (At2g13790) [42]. EFR signaling is not strongly inhibited in single *bak1* or *bkk1* null mutant. Only when using the hypomorphic allele *bak1–5*, which is strongly inhibited in PTI signaling, and the *bak1-5 bkk1-1* double mutant, a clear contribution of both SERK proteins to EFR signaling is detectable [41,42]. In Arabidopsis, the SERK family underwent an expansion and contains 5 members, which might have led to functional redundancy and diversification [78]. In rice, the SERK family contains only two members, OsSERK1 (Os08g07760) and OsSERK2 [54]. Only *OsSERK2*-silenced lines, but not *OsSERK1*-silenced lines are impaired in XA21-mediated immunity [54,79]. Rice expressing EFR and silenced for *OsSERK2* are impaired in elf18_*E.coli*_-triggered signaling. This suggests that rice SERK2 is the functional ortholog of Arabidopsis SERK3/BAK1 and SERK4/BKK1. Curiously, OsSERK2 is phylogenetically more closely related to Arabidopsis SERK1 (At1g71830) and SERK2 (At1g34210) [54]. Single mutants of Arabidopsis *serk1* and *serk2* in Arabidopsis are not impaired in elf18-triggered signaling, indicating that they do not play a role in the responses tested despite forming a ligand-induced complex with EFR [42]. The recruitment of phylogenetically distinct SERK proteins into the EFR plasma membrane signaling complex when comparing rice and Arabidopsis might also explain the different kinetics observed for the interaction of OsSERK2 and EFR in rice and BAK1 and EFR in Arabidopsis. In rice, OsSERK2 and EFR form ligand independent heteromers that do not appear to change within 15 min of ligand treatment (Figure 5B). In Arabidopsis, the interaction between SERK3 and EFR is only observable after ligand treatment within seconds to minutes [41,42].

EFR also directly interacts in a kinase activity independent manner with XB24, an enzymatically active ATPase. EFR rice lines overexpressing XB24 were slightly impaired in elf18_*E.coli*_-mediated defense signaling, especially at later time points (Figure 6E to 6H). XB24 therefore appears to be a negative regulator of EFR signaling in rice similar to its involvement in XA21-mediated immunity [53]. It remains to be determined if XB24 also enhances the autophosphorylation activity of EFR and employs identical mechanisms of regulation of EFR signaling as it does for XA21 signaling in rice.

A related study by Zipfel and colleagues shows that Arabidopsis XB24 also interacts with EFR [80]. Yet Arabidopsis *xb24* mutants are not impaired in elf18-triggered signaling. This might be due to the fact that Arabidopsis XB24 lacks several amino acids important for ATPase activity [80].

### EFR and EFR::XA21 confer quantitatively enhanced resistance to two weakly virulent *Xoo* isolates

EFR and EFR::XA21 rice plants are fully able to recognize the elf18 sequence derived from EF-Tu of *E. coli* in fully mature leaf tissue at sub-nanomolar concentrations (Figure 1 to 3, Table 1). Sequence analysis of over 20 *Xoo* isolates (Supplementary Table 2) revealed that the elf18 sequence in *Xoo* is highly conserved and contains two single amino acid changes at the second and fourth position when compared with elf18*_E.coli_* sequence (Supplementary Figure 8). The resulting elf18*_Xoo_* sequence was previously shown to be as active as elf18*_E.coli_* with an EC50 value of ~200 pM when used as double alanine substitution control in the medium alkalization assay of Arabidopsis cell cultures [27]. This observation also holds true for rice plants expressing EFR and EFR::XA21, because elf18*_E.coli_* and elf18*_Xoo_* elicited similar defense responses in these plants (Figure 3, Table 1). These observations led us to hypothesize that EF-Tu from *Xoo* would be fully recognized by EFR and EFR::XA21 expressing rice plants. Full length EF-Tu*_Xoo_* protein is most likely also readily available for recognition at the infection site of the xylem pathogen *Xoo* (Supplementary Figure 9) [68]. Based on this observation, we hypothesized that EFR, and especially EFR::XA21, expressing rice plants would be more resistant to Xoo. In the initial infection experiments, we used our fully virulent *Xoo* isolate PXO99A, to which rice lines expressing the rice immune receptor XA21 are fully resistant (Table 2) [28]. EFR-expressing plants were slightly more resistant to the fully virulent *Xoo* isolate PXO99A in 5 out of 8 infection assays (Figure 4A, Table 2). This partial disease resistant phenotype of EFR expressing rice plants is similar to the contribution of EFR to the resistance against the fully virulent *Pseudomonas syringae* pv. *tomato (Pto)* DC3000 isolate on Arabidopsis [35]. Several *Pto* DC3000 effectors have been shown to suppress EFR signaling during the infection process including AvrPto and HopAO1 [43,81]. *Xoo* PXO99A does not encode orthologs for any of these effectors but might be able to secrete *Xoo* specific effectors into rice cells to suppress PTI signaling initiated by EFR and other endogenous rice PRRs.

A screen of our diverse collection of *Xoo* isolates identified several *Xoo* isolates that are less virulent on Kitaake plants (Supplementary Table 4). EFR expressing rice plants show an enhanced resistance to the weakly virulent *Xoo* isolate MXO90 in 6 out of 6 experiments and to isolate *Xoo* NXO256 in 5 out 6 experiments for (Figure 4, Table 2). This quantitatively enhanced resistance phenotype in *Ubi::EFR::GFP* plants requires the expression of full-length EFR, because *Ubi::EFR::GFP-1*, which does not express EFR to any detectable level when measured by qRT-PCR and western blot analyses (Figure 1A and B), is not responsive to exogenous elf18_*E.coli*_ application (Figure 1C and B) and does not show an enhanced resistance phenotype for any *Xoo* isolate tested (Figure 4B, Table 2). Therefore, it is very likely that the recognition of EF-Tu from *Xoo* by EFR during the infection process leads to the quantitatively enhanced resistance phenotype of *Ubi::EFR::GFP* plants expressing EFR (Figure 4B, Table 2). This again is similar to the observation in Arabidopsis where the contribution of EFR towards disease resistance is more readily accessible when using hypo-virulent isolates of *Pto* such as *Pto* ΔCor^-^ and *Pto* ΔAvrPto/ΔAvrPtoB [35]. This is in contrast to previous gain-of-function experiments showing that the transgenic expression of EFR in tomato and *N. benthamiana* leads to a strong resistance response to taxonomically diverse bacterial pathogens under laboratory conditions [10]. The strength of the immune response conferred by the expression of EFR might be defined by the specific plant pathogen interaction and the infection methods used. We cannot completely rule out the possibility that the expression of EFR in rice does not lead to a fully functional immune receptor, because EFR might not interact appropriately with all required downstream immune signaling components. This defect in full signaling capacity might be the reason why the expression of EFR does not confer a qualitative resistance response to *Xoo* in rice. However, this is unlikely as the expression of EFR confers sensitivity to elf18*_Xoo_* (Figure 1–3, Table 1) similar to that observed for elf18_*E.coli*_ in Arabidopsis [26,27].

We also attempted to assess the resistance phenotype of *Ubi::EFR::XA21::GFP* plants, which was extremely difficult due to the observed early senescence phenotype at the infection stage using 6-week-old plants (Supplementary Figure 9). Nonetheless, we were able to perform two full experiments in which we infected fully mature leaves of *Ubi::EFR::XA21::GFP* plants that did not show any necrosis or early senescence phenotypes. In these experiments, *Ubi::EFR::XA21::GFP* plants were slightly more resistant to NXO256 in 1 out of 2 experiments, but not to MXO90 and PXO99A, however, they were less resistant than *Ubi::EFR::GFP* plants (Table 2). Although we did not perform as many experiments with the *Ubi::EFR::XA21::GFP* lines as we did with *Ubi::EFR::GFP* lines, it was clearly evident that the EFR::XA21 chimera receptor does not confer robust resistance to *Xoo* infection, despite its ability to detect and respond to elf18*_Xoo_* at sub-namolar concentrations (Figure 3, Table 1). These results are somewhat surprising because we and others previously hypothesized that the XA21 intracellular kinase domain would define the strong disease resistance phenotype mediated by XA21 [82,83]. For example, Kishimito et al. demonstrated that the expression of the chimeric receptor CeBIP::XA21 in rice confers enhanced resistance to the fungal pathogen *Magnaporthe oryzae* by increasing chitin perception and signaling [84]. However in our experimental system, expression of EFR::XA21::GFP in rice is not sufficient to trigger robust resistant to *Xoo*. This might be caused by inadequate immune signaling activation by this chimeric receptor, as suggested for EFR (see Discussion above). Alternatively these results suggest that the extracellular XA21 LRR, or the to-date unidentified ligand of XA21, contribute significantly to the strong disease phenotype of *XA21* rice plants against the bacterial pathogen *Xoo*. In the future, it will be important to assess the disease phenotype of rice plants expressing the reverse chimera XA21::EFR, in which the extracellular domain of XA21 is fused to the intracellular kinase of EFR. The analyses of the *Ubi::XA21::EFR* genotype and the knowledge of the ligand of XA21 will enable us to assess the role of the extracellular LRR of XA21 towards the strong disease phenotype of *XA21* plants. If the expression of XA21::EFR confers robust resistance to *Xoo* it will demonstrate that the ligand and the ectodomain of PRRs plays a more important role for the disease resistance response than previously anticipated.

### Stacking of PRRS as potential strategy to improve broad-spectrum disease resistance

Rice is unable to recognize elf18_*E.coli*_ (Figure 1–3) [85], and only the expression of the Arabidopsis PRR EFR enables rice to sense elf18_*E.coli*_ (Figure 1–3). It was shown recently that rice is able to detect a distinct part of EF-Tu from the bacterial pathogen *Acidovorax avenae* isolate N1141, which is located in its central region (Lys176 to Gly225) [85]. The authors hypothesize that rice possesses an alternate EF-Tu PRR that binds to the central region of EF-Tu. However, this central region of EF-Tu has only 66 % amino acid identity between *A. avenae* and *Xoo* and it is therefore difficult to speculate whether the endogenous EF-Tu receptor of rice would recognize EF-Tu from Xoo. It is currently unknown if Kitaake and other rice varieties such as the japonica rice cultivar Nipponbare are also able to sense this central region of EF-Tu. Generating rice with two independent EF-Tu immune receptors, EFR and the endogenous unknown PRR, would restrict the pathogens ability to mutate both recognition sites on the same protein concomitantly. It is therefore likely that the resistance mediated by both receptors would be more durable than by each single receptor. In the future, it will be important to test if transgenic rice plants expressing EFR provide resistance to *Xoo* or other bacterial pathogens such as X. *oryzae* pv. *oryzicola* under field conditions. The recognition of elf18*_Xoo_* by EFR during the initial low dosage *Xoo* infection through hydathodes and natural openings may be useful for limiting pathogen spread in the field especially in the presence of a second independent endogenous EF-Tu receptor recognizing another epitope. This stacking of several PRRs that recognize either different moieties of the same highly conserved protein or different PAMPs might be a valuable strategy to generate long-lasting broad-spectrum resistance [9]. Several recent studies, which describe the transfer of PRRs between different plants species, suggest that interspecies transfer of PRRs is feasible [10–13]. Field studies are needed to assess the full potential of this promising approach of increasing disease resistance by stacking multiple PRRs.

During the review process of this manuscript three newly published manuscripts also describe the successful transfer of EFR to a monocot crop, or reciprocally of *Xa21* to dicots. Ridout and colleagues demonstrate that the transgenic expression of EFR in wheat confers elf18_*E.coli*_ responsiveness and resistance to *Pseudomonas syringae* pv. *oryzae* (Schoonbeek et al., in press). Additionally, transgenic expression of EFR driven by the 35S promoter in the rice variety Zhonghua 17 conferred responsiveness to elf18_*E.coli*_, slight enhanced resistance to the bacterial pathogen *Acidovorax avenae* subsp. *avenae* at the seedlings stage but did not alter the immune response to *Xanthomonas oryzae* pv. *oryzae* PXO99A [61]. Conversely, transgenic expression of XA21 and EFR:XA21 in *Arabidopsis thaliana*, led to increased resistance to weakly virulent strain *Pseudomonas syringae* pv. *tomato* DC3000 COR^-^ and in the case of EFR::XA21 also towards *Agrobacterium tumefaciens* [80]. Taken together, all four publications highlight the interest in and the applicability of inter-class transfers of EFR and XA21, and potentially other plant PRRs, from dicots to monocots, and vice-versa.

## Methods

### Plant material and methods

Rice seeds were germinated in water-soaked filter paper for 5-7 days at 28°C and then transplanted into either 4.4-liter pots for plant inoculation assays and growth assessment or 3.5 liter pots for all other experiments. Plants were grown in an 80/20 (sand/peat) soil mixture in an environmentally-controlled greenhouse with temperature set to ~28-30°C and humidity to 75-85%. During winter months (November-April) artificial light supplementation was applied to obtain a day/night regime of 14/10.

### Generation of transgenic plants

Transgenic plants were generated as described previously [86]. Briefly, *pC::UBI::EFR::GFP* and *pC::UBI::EFR::XA21::GFP* were transformed into Kitaake calli by *Agrobacterium*-mediated transformation. Regenerated plants were selected on hygromycin. The presence of the transgene was confirmed in the T_0_ and each following generation by PCR using transgene specific primers (Supplementary Table 4).

### Rice crosses and progeny analysis

The confirmed T_2_ plants of *Ubi::EFR::GFP-9-4* were crossed to homozygous *OsSERK2RNA/* line *X-B-4-2* [54] or homozygous *XB24* overexpressor line *A109-6-5-1* [53]. In these crosses *Ubi::EFR::GFP* was used as pollen donor (male). Successful crosses were confirmed in the F1 generation and double transgenic plants were selected in the F2 generation by PCR reactions using specific primers for each transgene (Supplementary Table 4).

### Plasmid construction

We generated two plasmids for plant transformation using the *pNC1300* vector for final plant transformation [87]. The chimeric construct *EFR::GFP* and *EFR::XA21::GFP* in the *pENTR-D/TOPO* vector (Invitrogen) was generated as follows. We amplified two DNA fragments with about 25bp overlap using Phusion polymerase (Thermo). For the full *EFR* coding sequence we used primer combination of EFF-F and EFR_NOSTR on *EFR* CDS containing vector [35] and the 3’ GFP fusion part the primer combination GFPoverEFRF and GFPSTR on *pNC1300::UBI::Xa21::GFP* [88]. For the 5’ *EFR* fragment we used primer combination EFR-F and EFRectR on *EFR* CDS containing vector [35] and for the 3’ *XA21::GFP* fragment we used primer combination XaTMoverEFRF and GFPSTR on *pNC1300::UBI::Xa21::GFP* [88] (Supplementary Table 4). PCR products of the expected size were gel purified and 2 ul of each purified PCR product combined for a chimeric PCR reaction without primers using the following conditions: Denaturation 95°C for 1 min, Annealing 42°C for 30 seconds, Extension 72°C for 30 sec/kb, 12 cycles. The chimeric PCR reaction was diluted 1:1000 and used as template in a PCR reaction using the flanking primer combination EFR-F and GFPSTR for both chimeric constructs (Supplementary Table 4). PCR products of the expected size were gel purified and cloned into *pENTR-D/TOPO* vector (Invitrogen). The sequences of the chimeric genes *EFR::GFP* and *EFR::XA21::GFP* were confirmed by standard Sanger sequencing. Both *EFR::GFP* and *EFR::XA21::GFP* were flipped into the *pNC1300::UBI* transfer vector [87] by LRII clonase reactions (Invitrogen). Recombination reactions were confirmed by restriction analysis on the final vectors *pUbi::EFR::GFP* and *pUbi::EFR::XA21::GFP.*

For yeast-two hybrid assays we cloned the intracellular domain of EFR into *pLexA.* The intracellular domain of EFR was cloned into *pENTR-D/TOP0* vector (Invitrogen) using the primer combination EFR_2037_GW and EFR_stop_R on *pUbi::EFR::GFP*. We verified DNA sequence by standard Sanger sequencing. We also generated a clone where the aspartate (EFR849) in the catalytic loop of EFR was mutated to an asparagine in order to disrupt kinase activity [41]. The underlying point mutation was introduced by targeted point mutagenesis using the primer combination EFR_D-N_F and EFR_D-N_R on EFR ID in *pENTR-D/TOP0* (see above) using PCR conditions described previously [41]. We verified DNA sequence by standard Sanger sequencing. Both EFR ID and EFR (D849N) ID were flipped into the *pLexA* vector by LRII clonase reaction (Invitrogen). Recombination reactions were confirmed by restriction analysis on the final vectors *pLexA-EFR-ID* and *pLexA-EFR(D849N)-ID*.

For transient expression assays in *N. benthamiana*, we cloned the full coding sequence of XB24 without stop codon into pGWB11 [89]. XB24 without stop codon was cloned into pENTR-D/TOPO vector (Invitrogen) using the primer combination XB24_GW_F and XB24_w/o_stop on cDNA. We verified DNA sequence by standard Sanger sequencing. The CDS of XB24 without stop codon was flipped into pGWB11 by LRII clonase reaction (Invitrogen). Recombination reactions were confirmed by restriction analysis and sequencing of the final vector pGWB11-XB24.

### Agrobacterium-Mediated transient expression and immunoprecipitation from *N. Benthamiana*

Transient expression of XB24-FLAG from pGWB11-XB24 and EFR:::GFP from pEG101-EFR followed by immunoprecipitation were performed as previously described [41,80].

### Yeast two-hybrid assays

Yeast two-hybrid assays were performed as described previously [54,56] using the Matchmaker LexA two-hybrid system (Clontech). Yeast pEGY48/p8op-lacZ (Clontech) was co-transformed with *pLexA* and *pB42AD* vectors containing the indicated inserts by using the Frozen-EZ yeast transformation II kit (Zymo Research).

### Rice leaf tissue treatment with elicitors

Rice leaf tissue was treated with elicitors as described previously [54]. Leaves of 4-week-old greenhouse grown rice plants were cut into 2 cm long strips and incubated for at least 12 hours in ddH20 to reduce residual wound signal. Leaf strips were treated with water, 1 μM flg*22*_Pst_ peptide, purchased from Pacific Immunology, 500 nM elf18 peptides, purchased from Gene Script, or 50 μg/mL chitin, purchased from Sigma, for the indicated time. Leaf tissue was snap-frozen in liquid nitrogen and processed appropriately.

### qRT-PCR

Total RNA was isolated from rice plant tissues using TRIzol (Invitrogen), following the manufacturer’s protocol. Total RNA was treated with Turbo DNA-free DNAse (Ambion). RNA integrity was confirmed by standard agarose electrophorese in the presence of 0.1% SDS. 2 μg of total RNA was used for cDNA synthesis using the Reverse Transcriptase Kit (Applied Bio Science). Quantitative real time PCR (qRT-PCR) was performed on a Bio-Rad CFX96 Real-Time System coupled to a C1000 Thermal Cycler (Bio-Rad). For qRT-PCR reactions, the Bio-Rad SsoFast EvaGreen Supermix was used. qRT-PCR primer pairs used were as follows: Os04g10010-Q1/-Q2(5’-AAATGATTTGGGACCAGTCG-3’/5’-GATGGAATGTCCTCGCAAAC-3’) for Os04g10010 gene, PR10b-Q1/-Q2 (5’-GTCGCGGTGTCGGTGGAGAG-3’, 5’-ACGGCGTCGATGAATCCGGC-3’) for PR10b, EFR_ecto-Q1/-Q2 (5’-TGCATCTTTGCTCAAGCCAGGT-3’, 5’-GCGGCCACATGTGACTCCAA-3’) for EFR_ectodomain, Actin-Q1/-Q2 (5’-TCGGCTCTGAATGTACCTCCTA-3’/ 5’-CACTTGAGTAAAGACTGTCACTTG-3’) for the reference gene actin. qRT-PCR reactions were run for 40 cycles with annealing and amplification at 62°C for 5 sec and denaturation at 95°C for 5 sec. The expression levels of *Os04g10010, PR10b* and *EFR-ectodomain* were normalized to the actin gene expression level.

### Bacterial infection assays

To prepare *Xanthomonas oryzae* pv. *oryzae (XOO)* inoculum, *Xoo* isolates were spread-plated on peptone sucrose agar plates for 3 days, then washed off with water and adjusted to an OD_600_ of ~0.5, which corresponds to 5x10^8^ CFU/mL. Greenhouse-grown plants were transported into controlled growth chambers at the 5-to 6-week-old stage. Chamber conditions were set to ~28°C, 85% humidity and 14/10 day/night regime. Plants were allowed to acclimate to the chamber conditions for 2-3 days before being clip-inoculated with the *Xoo* inoculum[28]. In each plant 5–6 tillers were inoculated and in each tiller the two most recent fully developed leaves were clipped about 2 cm from the tip with scissors dipped in the *Xoo* inoculum. For each treatment 2-4 plants were inoculated yielding 20-40 inoculated leaves per treatment. Plants were incubated for 12–14 days post inoculation before disease lesions were scored.

*In planta* bacterial growth curves were performed as previously described [77].

### Rice biomass and yield assessment

Rice plants were grown as described above, with two seedlings in each 4.4-liter pot. At maturity, irrigation was stopped and plants were dried. Then total dry biomass and grain yield was weighted.

## MAP kinase assays

MAP kinase assays protocols were adapted from Arabidopsis [41]. Rice leave were ground to fine powder in liquid nitrogen and solubilised in better lacus buffer [50 mM Tris-HCl pH 7.5; 100 mM NaCl; 15 mM EGTA; 10 mM MgCl_2_; 1 mM NaF; 1 mM Na_2_MoO_4_.2H_2_O; 0.5 mM NaVO_3_; 30 mM β-glycerophosphate; 0.1% IGEPAL CA 630; 100 nM calyculin A (CST); 0.5mM PMSF; 1 % protease inhibitor cocktail (Sigma, P9599)]. The extracts were centrifuged at 16,000xg, the supernatant cleared by filtering through Miracloth and 5xSDS loading buffer added. 60 μg of total protein was separated by SDS-PAGE and blotted onto PVDF membrane (Biorad). Immunoblots were blocked in 5% (w/v) BSA (Fischer) in TBS-Tween (0.1%) for 1-2 H. The activated MAP kinases were detected using anti-p42/44 MAPK primary antibodies (1:1000, Cell Signaling Technology) overnight, followed by anti-rabbit-HRP conjugated secondary antibodies (Sigma).

## ROS production measurement in rice

Leaves of 3-to 4-week-old rice plants were cut longitudinal along the mid vein and then transverse into 1 to 1.5 mm thick leaf pieces. These leaf pieces were floated on autoclaved water overnight. The next morning two leaf pieces each were transferred into one well of a 96-well white plate containing 100 μl elicitation solution (20 µM LO-12 [Wako, Japan], 2 μg/ml HRP [Sigma]). Leaf pieces were treated with the indicated concentration of elf1*8_E.coli_* or elf18*_Xoo_* and time. ROS production was measured for 0.5s per reading with a high sensitivity plate reader (TriStar, Berthold, Germany).

## Western blot analysis

Total protein extracts from yeast, *Xoo* and rice plants and Western blot analyses were performed as previously described [41,54]. The primary antibodies used were as follows: Anti-OsSERK2 for detection of OsSERK2 [54], anti-GFP (Santa Cruze Biotech) for detection of EFR::GFP and EFR::XA21::GFP, anti-LexA (Clontech) for detection of LexA-fused proteins expressed in yeast from pLexA, anti-HA (Covance) for detection of HA-tagged proteins expressed in yeast from pB42AD and anti-EF-Tu (Thermo Fisher Scientific, PA5-27512) antibody to detect EF-Tu in *Xoo* protein preparations. The appropriate secondary antibody, anti-mouse (Santa Cruz Biotech) and anti-rabbit (GE Healthcare) coupled to horseradish peroxidase were used in combination with chemiluminescence substrates (Thermo) to detect proteins by exposure to film.

## Protein extraction and immunoprecipitation from rice tissue

Detached rice leaves from 4-week-old *Ubi::EFR::GFP, Ubi::EFR::XA21::GFP* or Kit plants were treated as described in *Rice leaf tissue treatment.* About 40mg of total protein in rice IP buffer (20mM Sodium Phosphate buffer pH 7.2, 150mM NaCl, 2mM EDTA, 10% Glycerol, 10mM DTT, 1% IGEPAL CA-630, plant protease inhibitor (Sigma P9599), 1mM PMSF, general protease inhibitor SigmaFast (Sigma), 1% PVPP) was used in combination with approximately 80 μL anti-GFP agarose slurry (Chromatek) for immunoprecipitation following the method described previously [41,42]. Immunoprecipitates were eluted from agarose beads by addition of 50-100 μL of 2xSDS loading buffer and heated to 70°C for 10 min. At least 50% of the eluate was loaded in order to detect anti-OsSERK2 in the immunoprecipitates.

## EF-Tu sequencing and alignment

*Xoo* carry two copies of the *tuf* gene that are 100% identical based on amino-acid sequence of the sequenced isolate PXO99A. The two *tuf* copies, PXO_04538 (copy 1) and PXO_04524 (copy 2), 1191 bp-long each, are separated in the PXO99A genome by 18,580 bp. We used the following primer sets to amplify the whole EF-Tu sequence of both copies. Copy 1: EF-Tu-F1: CCTTTCGTGAGCACCATTGC and EF-Tu-R4: AGCACGTAGACTTCGGCTTC; and copy 2: EF-Tu2-F: CCAAGAAGGGCTGAGTTCGT and EF-Tu-R2: CCTTGAAGAACGGGGTATGA. In both primer sets the forward prime anneal several bp upstream of the *tuf* gene to allow sequencing of the gene from the beginning without cloning. Phusion high-fidelity DNA polymerase (NEB) was used to PCR-amplify the *tuf* gene (~1300 bp). PCR amplicons were gel-purified and directly sequenced with the same forward and reverse primes.

## Statistical analysis and EC50 value determination

For statistical analysis of inoculated rice we used either Student’s t-test, Dunnet test or Tukey test depending on experimental set up, and as indicated in each experiment, using the JMP software.

ROS production dose response curves were modeled on the non-linear logistic 4p formula using the JMP software. EC50 were calculated at half maximal ROS production using the best fitted model.

## RNA isolation and quality assessment for RNAseq

Rice leaf strips of ~1.5 cm were collected from greenhouse grown, 4.5-week-old Kitaake and *Ubi::EFR::XA21::GFP-3-4* plants. After 12h of equilibration on sterile water, RNA was isolated from leaf strip tissue using the Spectrum™ Plant Total RNA Kit from Sigma-Aldrich and on-column DNAse treated to remove genomic DNA contamination following the manufacturer’s instructions. RNA was quantified using the Quant-IT™ Ribogreen® RNA Assay Kit. RNA quality was assessed on an Agilent Technologies Bioanalyzer.

## Sequencing

Stranded RNA-seq libraries were generated using the Truseq Stranded mRNA sample preparation kit (Illumina). mRNA was purified from 1 µg of total RNA using magnetic beads containing poly-T oligos. mRNA was fragmented using divalent cations and high temperature. The fragmented RNA was reversed transcribed using random hexamers and SSII (Invitrogen) followed by second strand synthesis. The double stranded cDNA was treated with end-repair, A-tailing, adapter ligation, and 10 cycles of PCR amplification. qRT-PCR was used to determine the concentration of the libraries. Libraries were sequenced on the Illumina Hiseq 2×150 bp.

## Gene expression analysis

Reads were aligned to reference genome (Osativa_MSU_v7) using TopHat version 2.0.7 [90,91]. Gene annotations (Osativa_MSU_v7.0) along with the EFR::XA21::GFP sequence were used for expression analysis. Sample correlation between Kitaake and *Ubi::EFR::XA21::GFP* replicates was performed with the R software using pearson correlation analysis of raw count data and plotted using the ggplot2 package [92,93]. Differential gene expression between Kitaake and *Ubi::EFR::XA21::GFP* was assessed using the Bioconductor edgeR package for R [94,95]. Gene ontology analysis was performed with the agriGO gene ontology tool using the *Oryza sativa* dataset reference (http://bioinfo.cau.edu.cn/agriGO/).

## *Xoo* supernatant preparation

*Xoo* cultures were grown as described before [77]. In short, cells were grown in 10 mL of yeast extract broth (YEB) media (5 g/L yeast extract, 10 g/L tryptone, 5 g/L NaCl, 5 g/L sucrose, 0.5 g/L MgSO_4_, pH 7.3) to an OD600 of ~1.5, spun down and resuspended in 2 mL of M9 minimal media containing 1.5% glucose and 0.3% casamino acids. Cultures were further incubated at 28°C for 48 h. Before harvest a sample of total cells was collected and then the cells were spun down and the supernatant was passed through a 0.22 μM-filtering unit, representing the secreted fraction.

For mass-spectrometry (MS) analysis PXO99 cells were grown in M9 media until OD_600_ of ~ 0.150. Cells were spun down at 10,000xg for 15 min and the supernatant was collected and filtered through a 0.22 μM filter. For mass-spectrometry (MS) analysis PXO99 cells were grown in M9 media until OD_600_ of ~ 0.150. Cells were spun down at 10,000xg for 15 min and the supernatant (> 50 ml) was collected and filtered through a 0.22 μM filter. Four times volume of ice-cold acetone was added to the supernatant sample, vortexed vigorously and incubated at −20 °C for 6 hours with occasional agitation. Samples were then spun down at 15,000xg for 10 min. Residual acetone was air dried to evaporate from the protein pellet, after which proteins were resuspended in 50 mM Tris, 8 M Urea (pH 9.0) and quantified using the BCA assay (Biorad). Samples were then reduced (10 mM DTT; 30 min), alkylated (50 mM IAA; 20 min), and subjected to 4x sample volume dilution using 50% methanol to reduce Urea concentration. Samples were next digested overnight at room temperature using Trypsin (Promega Mass Spec grade) at 1:10 enzyme to protein ratio. Speedvac digested peptides were then resuspended in Buffer A (80% ACN; 0.1% TFA) and desalted using C18 Micro SpinColumn (Harvard Apparatus).

## Identification of proteins by LC-MS/MS

The digested secretome samples were analyzed on an Agilent 6550 iFunnel Q-TOF mass spectrometer (Agilent Technologies) coupled to an Agilent 1290 LC system (Agilent). The desalted peptide samples (40 μg) were loaded onto a Ascentis Peptides ES-C18 column (2.1 mm × 100 mm, 2.7 μm particle size; Sigma-Aldrich) via an Infinity Autosampler (Agilent Technologies) with buffer A (2% acetonitrile, 0.1% formic acid) flowing at 0.400 ml/min. The column compartment was set at 60°C. Peptides were eluted into the mass spectrometer via a gradient with initial starting conditions of 5% buffer B (98% acetonitrile, 0. 1% formic acid) increasing to 30% buffer B over 30 minutes, then to 50% buffer B in 5 minutes. Subsequently, buffer B concentration was increased to 90% over 1 minute and held for 7 minutes at a flow rate of 0.6 mL/min followed by a ramp back down to 5% buffer B over one minute, where it was held for 6 minutes to reequilibrate the column. Peptides were introduced to the mass spectrometer from the LC via a Dual Agilent Jet Stream ESI source operating in positive-ion mode. A second nebulizer was utilized for the introduction of reference masses for optimal mass accuracy. Source parameters employed Gas Temp (250°C), Drying Gas (14 L/min), Nebulizer (35 psig), Sheath Gas Temp (250°C), Sheath Gas Flow (11 L/min), VCap (3500 V), Fragmentor (180 V), OCT 1 RF Vpp (750 V). The data were acquired with the Agilent MassHunter Workstation Software, LC/MS Data Acquisition B.05.00 (Build 5.0.5042.2) operating in Auto MS/MS mode. A maximum of 20 precursors per cycle were selected for MS/MS analysis, limited by charge states 2, 3 and >3, within a 300 to 1400 m/z mass range and above a threshold of 1500 counts. The acquisition rate was set to 8 spectra/s. MS/MS spectra were collected with an Isolation Width at Medium (~4 m/z) resolution and collision energy dependent on the m/z to optimize fragmentation (3.6 × (m/z) / 100 – 4.8). MS/MS spectra were scanned from 70 to 1500 m/z and were acquired until 40000 total counts were collected or for a maximum accumulation time of 333 ms. Former parent ions were excluded for 0.1 minute following selection for MS/MS acquisition.

## MS/MS data analysis

The acquired data were exported as .mgf files using the Export as MGF function of the MassHunter Workstation Software, Qualitative Analysis (Version B.05.00 Build 5.0.519.13 Service Pack 1, Agilent Technologies) using the following settings: Peak Filters (MS/MS) the Absolute height (≥20 counts), Relative height (#x2265;0.100% of largest peak), Maximum number of peaks (300) by height; for Charge State (MS/MS) the Peak spacing tolerance (0.0025 m/z plus 7.0 ppm), Isotope model (peptides), Charge state Limit assigned to (5) maximum. Resultant data files were interrogated with the Mascot search engine version 2.3.02 (Matrix Science) with a peptide tolerance of ±50 ppm and MS/MS tolerance of ±0.1 Da; variable modifications Acetyl (N-term), Carbamidomethyl (C), Deamidated (NQ), Oxidation (M); up to one missed cleavage for trypsin; Peptide charge 2+, 3+ and 4+; and the instrument type was set to ESI-QUAD-TOF. Data was acquired and exported using MassHunter (Agilent Technologies) and resultant MS/MS data was analyzed using Mascot (Matrix Sciences) against a custom database comprising the RefSeq PXO99A proteins (ca. 13,500 proteins) and all Viridiplantae proteins (ca. 565,000 proteins) available through NCBI. Thresholds were also set to reduce the false discovery rate (p<0.05) and ensure significant peptide and protein matching. Protein and peptide matches identified after interrogation of MS/MS data by Mascot were filtered and validated using Scaffold (version 4.1.1, Proteome Software Inc., Portland, OR). Peptide identifications were accepted if they could be established at greater than 95.0% probability by the Peptide Prophet algorithm [96] with Scaffold delta-mass correction. Protein identifications were accepted if they could be established at greater than 99.0% probability and contained at least 1 identified peptide (at 95% and greater).

Statistical analysis was performed using the Tukey-Kramer HSD test. Different letters indicate significant differences (p < 0.05).

Statistical analysis was performed using the Tukey-Kramer HSD test. Different letters indicate significant differences (p < 0.05). These experiments were repeated at least three times with similar results.

## Acknowledgments

We thank Markus Albert for discussion and technical advice on setting up the ROS assay in rice. BS and OB thank UAW5810 UC postdoc Union for constantly improving UC post-doctoral scholar working and living conditions.

---

**Supplementary Figure 1: Phylogenetic tree of XA21 with its ten closest homologs in rice and Arabidopsis**

**Supplementary Figure 2: Schematic representation of PRR clones used in this study**

Numbers indicate amino acid residues of fusion points. Drawn to approximate scale.

**Supplementary Figure 3: Preliminary protein expression of analysis in EFR::GFP and EFR::XA21::GFP T_1_ plants** Western blot analysis of pooled total protein fractions of several PCR positive T_1_ plants for each independent T_0_ line of *Ubi::EFR::GFP* and *Ubi::EFR::XA21::GFP* transgenic rice plants.

**Supplementary Figure 4: Full anti-GFP western blot on total protein extracts from fully mature leaves of Kitaake,** *Ubi::EFR::GFP* **and** *Ubi::EFR::XA21::GFP* **lines**

**Supplementary Figure 5: MAP kinase activation in rice leaves treated with flg22 and chitin** Fully mature leaves of Kitaake were treated with (A) 1 μM flg22_Pta_ or (B) 50 μg/ml chitin for the indicated time. Upper panel anti-p42/44 MAP kinase western blot on total protein extracts, lower panel CBB stain of membrane as loading control.

**Supplementary Figure 6: Total ROS production over 3 hours in response to 100nM elf18_*E.coli*_ in rice plants expressing EFR::GFP and EFR::XA21::GFP, and Kitaake control**

Bars depict average relative light production ± SE of at least six biological replicates. Statistical analysis was performed using the Tukey-Kramer HSD test. Different letters indicate significant differences (p < 0.05). These experiments were repeated at least three times with similar results.

**Supplementary Figure 7: Dose response curves to elf18_*E.coli*_ and elf18*_Xoo_* in rice expressing EFR::GFP or EFR::XA21::GFP**

Individual points depict single measurements of total ROS production over 3 hours. The line depicts the best-fitted model to the non-linear logistic 4p formula using the JMP software package.

**Supplementary Figure 8: Alignment of the elongation factor-Tu (EF-Tu) protein sequences among *Xoo* isolates, *A. avenae* and*E. coli* as reference sequences.**

The elf18 sequence is marked with black line and the EF-Tu EFa50 region (176-225) is marked with a hatched line. The EF-Tu protein is present in all tested *Xoo* isolates in two copies. Sequence analysis of the first ~250 amino acid of both copies in 20 *Xoo*isolates revealed that they are 100 % identical therefore only one EF-Tu*_Xoo_* sequence is shown. The first 18 amino acids (elf18) of *Xoo* contain two base-pair substitutions at positions 2 and 4, as compared with the sequence of *E. coli.* The 176-225 (EFa50) region has 66 % identity between *Xoo* isolates and *A. avenae*, while the full-length protein has 83 % identity.

**Supplementary Figure 9: *Xoo* EF-Tu is detected in cell-free supernatants.** Western blot analysis with (A) an anti-EF-Tu antibody and (B) mass spectrometry analysis of PXO99 cell-free supernatants reveal that EF-Tu is present in the outer cellular space under *in vitro* growing conditions in rich media.

**Supplementary Figure 10: Transgenic expression of EFR in rice does not negatively impact growth or yield**. (A) Total dry weight (top) and total yield (bottom) analysis of Kitaake compared with two independent lines of *Ubi::EFR::GFP* and *Ubi::EFR::XA21::GFP* transgenic lines. Statistical analysis was done using the Tukey-kramer HSD test. Different letters indicate significant difference at the 0.05 alpha level. (B) Pictured illustration of Kitaake vs. *Ubi::EFR::GFP and Ubi::EFR::XA21::GFP* lines at the vegetative stage (top), and Kitaake Vs. *Ubi::EFR::GFP* at the flowering stage. Boxed with dashed line is a zoon-in image of a characteristic necrosis appearing in the *Ubi::EFR::XA21::GFP* line at 6-week stage.

**Supplementary Figure 11: Pairwise analysis of whole transcriptome profile of Kitaake and *Ubi::EFR::XA21::GFP* plants at the 4-week stage.** Heatmap and dendogram of Pearson’s correlation coefficients between Kitaake and *Ubi::EFR::XA21::GFP-3-4.* Pearson correlation coefficients were based on logarithmic scaled raw count data.

**Supplementary Figure 12: GO analysis of differentially regulated genes between Kitaake and *Ubi::EFR::XA21::GFP* at the 4-week stage.**

A, GO terms associated with differentially up-regulated genes between Kitaake and *Ubi::EFR::XA21::GFP-3-4* plants. No significant GO term enrichment observed between reference and up-regulated gene set. B, GO terms associated with differentially down-regulated genes. A significant portion of down-regulated genes is associated with oxidoreductase activity (p = 032, FDR = 0.042).

**Supplementary Figure 13: The transgenic expression of EFR in rice slightly inhibits bacterial replication of three different *Xoo* isolates at some time points.**

Rice lines were inoculated using the leaf-clipping method as described in materials and methods. Bacterial burden was recorded at 4 time points (0, 3, 8, 12 days post inoculation). Statistical analysis was done using t-test for each time point separately.

**Supplementary Figure 14: XA21 interacts with OsSERK2, XB3, XB15 and XB24 in the yeast-two hybrid assay.** Yeast two-hybrid assay between XA21K668 (668-1025aa) and OsSERK2 ID (260-628aa), XB3 full-length (FL) (1-450aa), XB15 FL (1-639aa) and XB24 FL (1-198aa). The blue color indicates nuclear interaction between the two co-expressed proteins.

**Supplementary Figure 15: All fusion proteins of EFR, EFR(D-N), OsSERK2, XB3, XB15, XB24 and GUS are expressed in yeast**. Anti-LexA (upper panel) and anti-HA (lower panel) western blot analysis on total yeast protein extracts from the yeast two-hybrid experiment shown in Figure 5A.

\# indicates full-length fusion protein LexA-EFR-ID and LexA-EFR(D-N)-ID

$ indicates full-length fusion protein LexA-GUS

% indicates full-length fusion protein AD-XB15

& indicates full-length fusion protein AD-OsSERK2-ID

@ indicates full-length fusion protein AD-XB24

* indicates full-length fusion protein AD-XB3

**Supplementary Figure 16: Stable expression of EFR in the F2 generation of double transgenic lines of a cross between *Ubi::EFR::GFP* and *OsSERK2RNAi* or *XB24OE***

*EFR* expression in double transgenic lines *Ubi::EFR::GFP × OsSERK2RNAi* (A) or *Ubi::EFR::GFP × XB240E* (B). Expression levels were measured by qRT-PCR and normalized to actin reference gene expression. Bars depict average expression level ± SE of three technical replicates.

**Supplementary Figure 17: Up-regulation of the two SA marker genes *NH1* and chalcone synthase (*CHS*) in 6-week old *Ubi::EFR::XA21::GFP* plants**

Expression of *CHS* (A) and *NH1* (B) in leaf tissue of 5-week-old *Ubi::EFR::XA21::GFP* plants. Expression levels were measured by qRT-PCR and normalized to actin reference gene expression. Bars depict average expression level ± SE of three technical replicates.

**Supplementary Table 1: T_1_ Segregation analysis of newly generated transgenic lines used in this study**

**Supplementary Table 2: Nomenclature and origin of *Xoo* isolates used in this study**

**Supplementary Table 3: Differentially expressed gene list**.

Differentially expressed genes between Kitaake and *Ubi::EFR::XA21::GFP-3-4.* Differentially expressed genes were selected using a false discovery rate of ≤ 0.05 and an absolute log fold change ≥ 2. Tables at the bottom of worksheets summarize gene ontology information from AgriGO.

**Supplementary Figure 4: Inoculation with different *Xoo* isolates.**

Rice inoculation with ten different *Xoo* isolates to identify weakly virulent *Xoo* isolates compared with the fully virulent isolates PXO99A.

**Supplementary Table 5: Table of primers used in this study**

